# A Nerve-Dependent NGF Receptor Switch Controls Corneal Epithelial Renewal In Neurotrophic Keratopathy

**DOI:** 10.64898/2026.06.30.735622

**Authors:** Arif Hussain, Kiana Tajdaran, Ramkumar Katturajan, Kaveh Mirmoeini, Jordan Crabtree, Alaa I. Quddam, Xiangbing Wu, Noam Blum, Daniel J. Konig, Jay S. Pepose, Asim Ali, Ruby Shalom-Feuerstein, Tessa Gordon, David R. Kaplan, Gregory H. Borschel, Konstantin Feinberg

**Affiliations:** Department of Surgery, Indiana University School of Medicine; Indianapolis, Indiana, USA; Program in Neurosciences and Mental Health, Hospital for Sick Children; Toronto, Ontario, Canada; Department of Genetics & Developmental Biology, The Rappaport Faculty of Medicine & Research Institute, Technion Integrated Cancer Center, Technion-Israel Institute of Technology; Haifa, Israel; Department of Ophthalmology and Visual Sciences, Washington University School of Medicine; St. Louis, MO, USA; Pepose Vision Institute; St. Louis, MO, USA; Department of Ophthalmology and Vision Sciences, University of Toronto and the Hospital for Sick Children; Toronto, Ontario, Canada; Departments of Medical Genetics, University of British Columbia; Vancouver, British Columbia, Canada; Department of Molecular Genetics, University of Toronto; Toronto, Ontario, Canada; Department of Ophthalmology, Eugene and Marilyn Glick Eye Institute, Indiana University School of Medicine; Indianapolis, Indiana, USA; Stark Neurosciences Research Institute, Indiana University School of Medicine; Indianapolis, Indiana, USA

## Abstract

Corneal epithelial integrity depends on continuous epithelial renewal by limbal epithelial stem cells (LESCs), a process tightly linked to sensory innervation. Loss or impairment of innervation causes neurotrophic keratopathy (NK), a sight-threatening degenerative disease for which rhNGF, the only FDA-approved pharmacologic therapy, often has limited efficacy in advanced or refractory disease. The mechanistic basis for this limited response remains unclear. Using surgical, genetic, and pharmacologic approaches in a rodent model of NK with corneal Schwann cell ablation or structural and functional denervation, together with primary human LESCs, we examined how denervation alters NGF receptor signaling during epithelial repair. In innervated corneas, NGF promoted epithelial regeneration through TrkA. Denervation, however, increased expression of a second NGF receptor, anti-regenerative p75^NTR^, and activation of its effector JNK, and reduced the activity of the TrkA effector AKT in LESCs. In this altered receptor context, denervation-induced elevation of endogenous NGF amplified p75^NTR^ signaling, thereby explaining the failure of topical rhNGF to rescue severely denervated NK phenotype corneas.

Conversely, selective TrkA activation, either with the clinical-stage agonist tavilermide, or pharmacologic or genetic inhibition/ablation of p75^NTR^, restored AKT signaling and rescued epithelial healing in denervated corneas independent of reinnervation. These findings identify a nerve-dependent NGF receptor switch as a key regulator of corneal epithelial renewal and establish receptor-selective modulation as a mechanistically rational therapeutic strategy for treating NK.

**One Sentence Summary:** Corneal denervation shifts limbal epithelial stem cell signaling from pro-regenerative TrkA–AKT toward anti-regenerative p75^NTR^–JNK, explaining the limited efficacy of recombinant human nerve growth factor (rhNGF; cenegermin), the only approved pharmacologic therapy for neurotrophic keratopathy, in severely denervated corneas and identifying receptor-selective modulation as a mechanistically distinct therapeutic strategy.

## INTRODUCTION

Corneal clarity is essential for vision. The corneal surface is maintained by continuous renewal of a stratified, non-keratinized epithelium derived from limbal epithelial stem cells (LESCs) ^1–3^. LESCs reside in the limbus, a narrow epithelial transition zone between the cornea and conjunctiva that constitutes the LESC niche ^4^. During epithelial maintenance and following injury, LESC progeny differentiate into transient amplifying cells (TACs) that migrate centripetally and differentiate into basal, wing and superficial epithelial cell layers, replenishing the corneal surface. Homeostatic epithelial renewal is sustained by long-term LESC self-renewal, whereas injury triggers rapid re-epithelialization through accelerated LESC activation and epithelial migration ^5–8^. Such rapid adaptation requires mechanistic responsiveness and flexibility.

The cornea is the most densely innervated tissue in the body, with sensory nerve fibers mediating pain sensation, tearing, and blink reflexes ^9^. Beyond their protective role, corneal nerves trophically regulate LESC maintenance and activity, contributing to both homeostatic and injury-induced epithelial renewal ^4,10^. In the limbus, nerve fibers run adjacent to LESCs and directly contact epithelial cells ^11,12^. Loss or impairment of corneal innervation, caused by infection, trauma, tumors, acquired or congenital neurological disorders, or metabolic conditions such as diabetes mellitus, causes neurotrophic keratopathy (NK), a rare but potentially blinding degenerative corneal disease characterized by impaired wound healing, epithelial breakdown, stromal thinning and melting, scarring, and progressive loss of corneal transparency ^13,14^. Among the trophic factors implicated in innervation-dependent corneal epithelial renewal ^15,16^, NGF has emerged as a central but incompletely mechanistically understood regulator ^17,18^. Recombinant human NGF (rhNGF, cenegermin, marketed as Oxervate by Dompé, Milan) is the only FDA approved pharmacological treatment for NK ^19,20^. Several studies suggest that NGF promotes epithelial healing indirectly by supporting corneal innervation, as evidenced by increased corneal sensitivity and subbasal nerve regeneration in cenegermin-treated patients with partial but not complete corneal denervation ^21–24^. In addition to its neurotrophic role ^25^, NGF exerts effects in non-neuronal tissues, including regulation of regenerative processes ^18,26–29^. NGF can exert trophic effects in a paracrine and autocrine manner in keratinocytes ^30^.

NGF signaling is mediated through two structurally and functionally distinct receptors, TrkA and p75^NTR^ ^25^. These receptors can act independently or cooperatively, yet often exhibit antagonistic functions with signaling outcomes determined by their activation state, relative expression, and cellular context ^25,26^. Notably, p75^NTR^ can act unliganded or be activated by multiple neurotrophic factors in their mature or precursor forms, including NGF and proNGF, whereas TrkA is preferentially activated by mature NGF, with NT-3 serving as a lower-affinity alternative ligand ^31–34^. p75^NTR^ can modulate TrkA signaling by altering ligand binding properties and receptor conformation, suggesting that it functions as a signaling rheostat when complexed with TrkA ^31–35^. While TrkA primarily mediates pro-survival, proliferative, and differentiation signals, p75^NTR^, a member of the TNF receptor superfamily, can promote cell cycle arrest, suppress proliferative signaling or promote apoptosis depending on cellular context and the availability of TrkA co-signaling ^36,37^.

In the cornea, NGF is predominantly expressed in the limbus and both TrkA and p75^NTR^ are expressed by basal epithelial cells, including LESCs ^17,38–40^. Therefore, in addition to its neurotrophic activity, NGF may directly regulate LESC function. However, whether NGF acts directly on LESCs to regulate epithelial renewal, and how NGF signaling affects LESC survival and activity, remain unresolved.

Consistent with this uncertainty, the clinical efficacy of rhNGF remains limited and variable. Cenegermin fails to induce complete healing and prevent recurrence in a substantial proportion of patients. The clinical response is frequently accompanied by restoration of corneal sensation and evidence of reinnervation in partially innervated corneas, whereas benefit is limited in anesthetic corneas ^24,41–43^, suggesting that therapeutic efficacy depends partly on residual innervation. In *ex vivo* and *in vitro* studies, exogenous NGF has been implicated in regulating both homeostatic and injury-induced epithelial renewal through effects on human LESCs ^17,18,39,44^. Consistently, loss-of-function approaches including pharmacological inhibition of TrkA or neutralization of endogenous NGF impaired LESC proliferation, colony formation and migration ^17,39,44^, suggesting a positive role for TrkA in maintenance and activity of LESCs. Notably, freshly isolated LESCs exhibit high p75^NTR^ and low TrkA expression, followed by a progressive decrease in p75^NTR^ and increase in TrkA levels during differentiation ^17^. Paradoxically, endogenous NGF levels increase in both corneal de-epithelialization associated with rapid and complete healing, and corneal denervation, which results in NK and complete failure of epithelial renewal ^12,18,45^. Moreover, NGF accelerates healing following experimental corneal de-epithelialization, while neutralization of endogenous NGF only modestly delays this process ^18^. Together, these observations raise fundamental questions: **a**) How does corneal innervation regulate NGF receptor signaling in LESCs to control epithelial renewal, and why does exogenous NGF fail to heal severely denervated corneas?, and **b**) If NGF levels rise in both healing and non-healing corneas, what receptor-level mechanism determines whether NGF signaling promotes or suppresses epithelial regeneration?

This study combines surgical, genetic, pharmacological, and *in vitro* approaches to define the NGF receptor signaling mechanism by which corneal innervation controls LESC activity and epithelial renewal. We observed that the balance between TrkA-AKT and p75^NTR^-JNK signaling in LESCs determines precise epithelial regenerative capacity. Corneal innervation maintains this balance through Schwann cell-dependent regulation of p75^NTR^ expression, and denervation inverts it thereby driving the NK phenotype through p75^NTR^ dominance that is amplified by elevated endogenous NGF. These findings not only provide a mechanistic explanation for the failure of topical rhNGF to rescue epithelial healing in severely denervated NK corneas, but also support selective TrkA activation or p75^NTR^ inhibition, as a mechanistically rational therapeutic strategy for NK that may complement or in some cases substitute for surgical corneal neurotization.

## RESULTS

### Topical rhNGF does not enhance epithelial healing or re-innervation in fully denervated corneas

Although NGF is clinically used to treat NK, its effect on corneal epithelial renewal has not been evaluated in a fully denervated NK model. Given the limited efficacy of rhNGF (cenegermin) in severe NK cases with near-complete or complete loss of corneal innervation ^46,47^, we examined its effect on epithelial healing in denervated corneas using our rodent NK model. In this model, unilateral corneal denervation was achieved by electrocautery of the ophthalmomaxillary branch of the trigeminal nerve ^12,48^.

Denervated rats received a single daily topical dose of 10 μl of rhNGF at 50 μg/ml — a concentration chosen to ensure receptor engagement while accounting for precorneal drug loss following topical instillation and the accelerated healing kinetics of the rodent cornea — beginning immediately after denervation (Fig. 1A).

**Figure 1.**
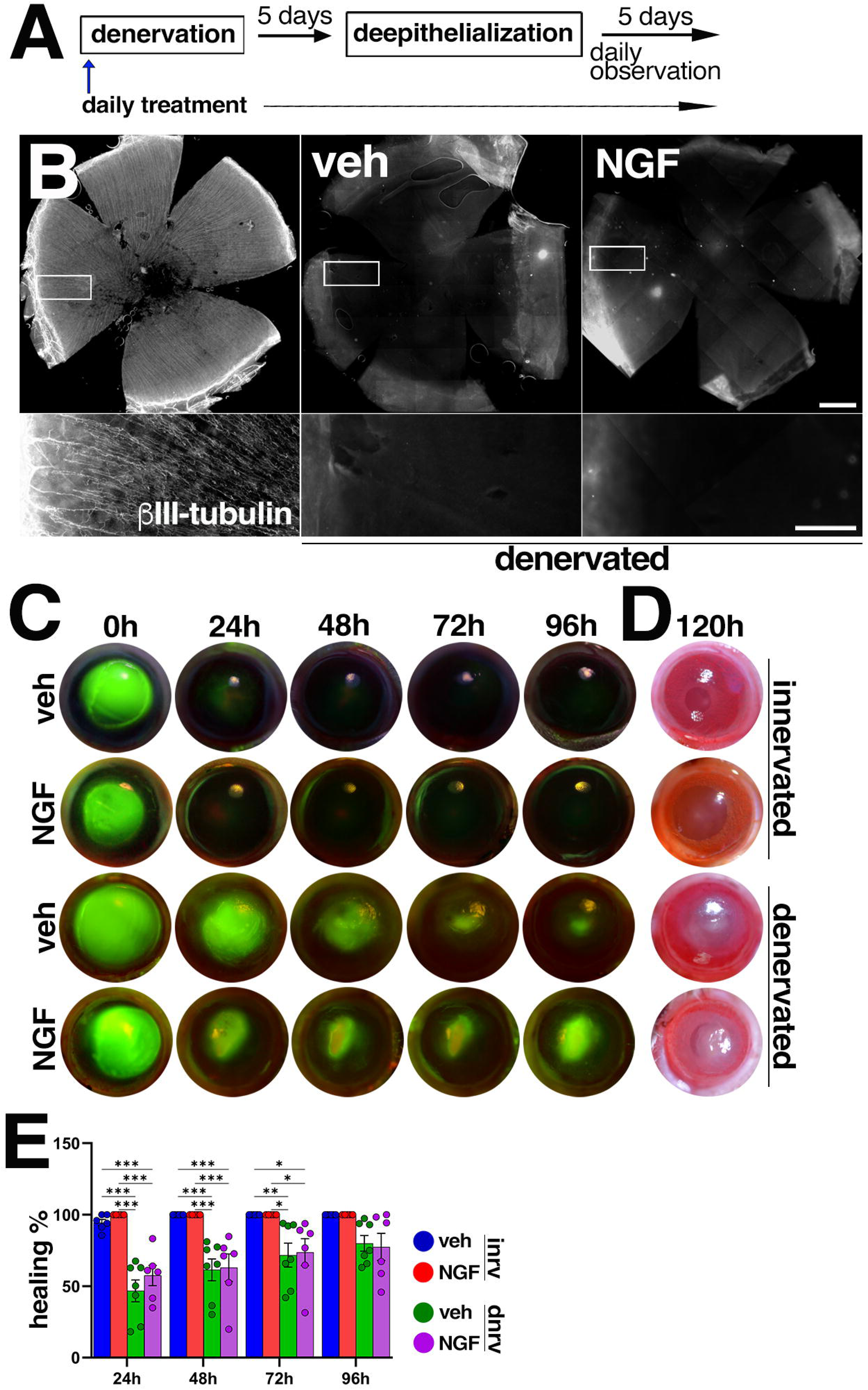
Topical NGF fails to induce epithelial healing of NK cornea. (**A**) Schematic description of the experiment in (C). (**B**) Representative fluorescent images of whole mount βIII-tubulin-immunolabeled normally innervated (left panel) or denervated (central and right panels) vehicle alone (“veh”) or rhNGF (NGF) treated rat corneas. Scale bars: upper panel - 1μm; lower (magnified enclosed image) panel – 400 µm. (**C**) Representative live photographs of fluorescein-stained rat normally innervated or denervated de-epithelialized vehicle only or rhNGF treated corneas during the indicated period. **0h** indicates corneal condition five days post denervation, immediately after de-epithelialization. (**D**) Representative live bright field images of the corneas as per (C), 120 hours post-de-epithelialization. (**E**) Quantitative representation of corneal healing as per (C) of innervated “inrv” or denervated “dnrv” corneas. Each data point represents the fluorescein-negative (healed) corneal area. *n* ≥ 5, ANOVA.

On day five post-denervation, corneal epithelial wounds were created using an Amoils brush, and healing was monitored for five days by fluorescein staining ^12,48^. Normally innervated rhNGF-treated and denervated vehicle-treated corneas served as positive and negative controls, respectively. Tarsorrhaphy (suturing of the eyelid margins) was performed throughout the experiment (and in all further presented corneal healing assays) to prevent secondary injury ^48^. In human patients, rhNGF-induced corneal healing is often accompanied by recovery of corneal sensation in partially denervated corneas ^20,47^. To confirm successful denervation and assess potential re-innervation, corneas were harvested prior to de-epithelialization and immunostained for the neuronal marker βIII tubulin. While dense limbal and epithelial innervation was observed in control corneas, no nerve fibers were detected in either rhNGF- or vehicle-treated denervated corneas (Fig. 1B), indicating that rhNGF did not promote re-innervation within this timeframe. As expected, complete epithelial healing occurred in normally innervated corneas treated with either vehicle or rhNGF. In accordance with previous reports ^12,48^, denervated vehicle-treated corneas exhibited partial wound closure accompanied by scarring and opacification, consistent with the NK phenotype (Fig. 1C–E) ^12,48^. Differing from the reported capsaicin-induced partial corneal denervation studies ^45^, yet consistent with clinical observations ^41–43^, rhNGF treatment failed to promote epithelial closure in denervated de-epithelialized corneas (Fig. 1C–E). Similar results were observed at higher dosing (0.5 mg/ml twice daily; not shown), indicating that the lack of efficacy was not attributable to insufficient NGF availability. Collectively, these results demonstrate that topical rhNGF neither restores epithelial healing nor induces re-innervation in fully denervated corneas.

### Modulation of NGF receptors affects the activity and maintenance of LESCs

Previous studies demonstrated expression of the NGF receptors TrkA and p75^NTR^ in limbal epithelial cells ^17,39,40,49^. Consistent with these findings, our immunohistochemical analysis of mouse (Fig. 2A–D) and rat (not shown) corneas confirmed expression of both receptors in limbal cells. Consistent with previous reports, p75^NTR^ immunostaining was predominantly associated with basal limbal epithelial ΔNp63-positive cells (ΔNp63 is a marker for these cells ^50^), whereas TrkA expression was detected in both ΔNp63-positive and ΔNp63-negative suprabasal limbal epithelial cells (Fig. 2A–D).

**Figure 2.**
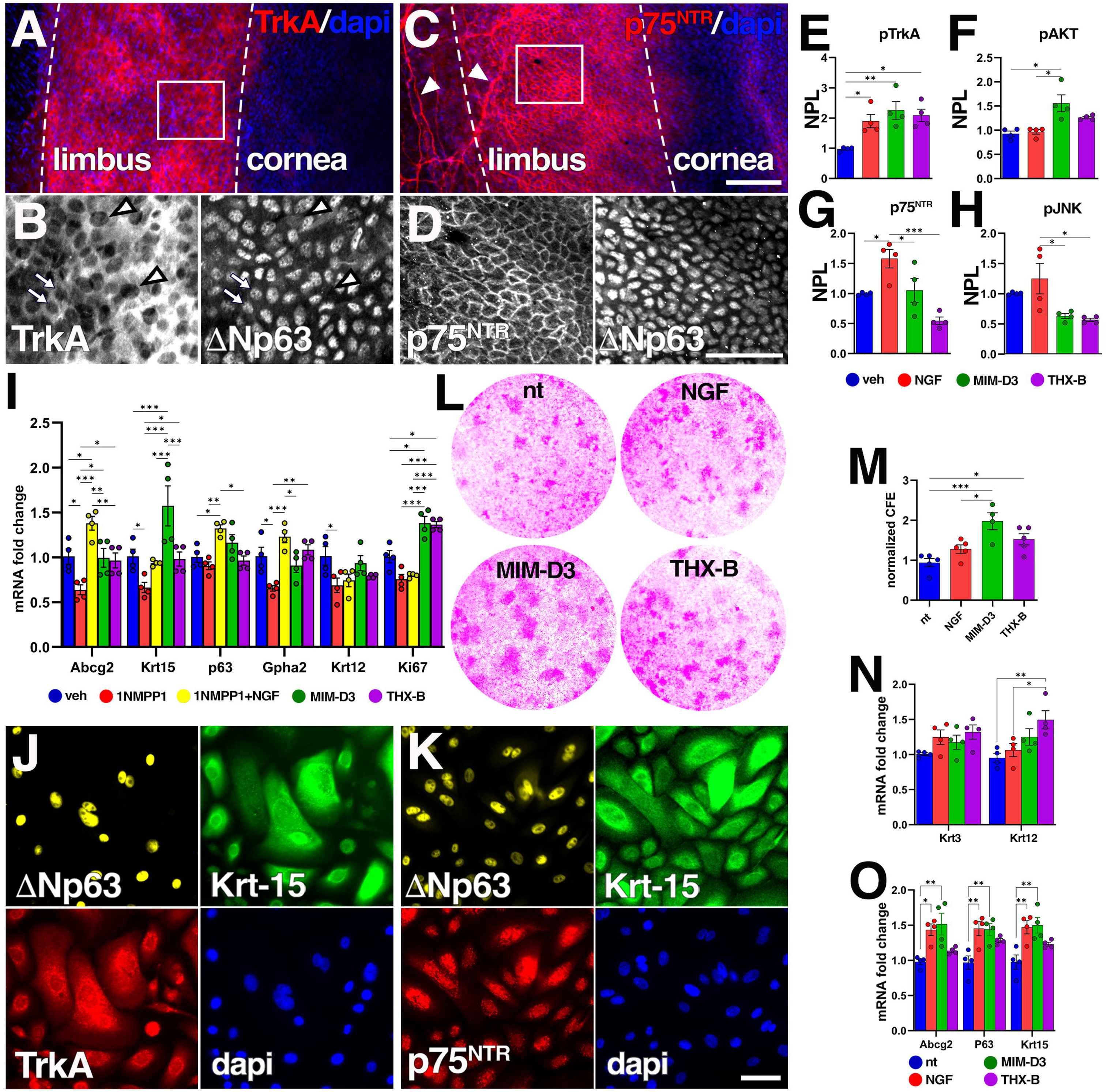
Modulation of the NGF receptors affects LESCs. (**A**-**D**) 15 μm extended depth-of-focus representative immunofluorescent images of mouse corneal epithelium demonstrate TrkA (red) in (A) or p75^NTR^ (red) in (C), and dapi double-positive cells at the denoted limbal area. (**B**), (**D**) magnified split images of the squared area in (A), (C), respectively, of co-immunolabeled for ΔNp63. In (C) arrowheads denote p75^NTR^ positive nerves-ensheathing Schwann cells. In (B) arrows denote TrkA/ΔNp63 double-positive subepithelial cells and arrowheads denote larger ΔNp63-negative epithelial cells. Scale bars, [(A), (C)] 100 μm; [(B), (D)] 50 μm. (**E-H**) Quantitative representation of normalized protein level (NPL) as per Fig. S1A. *n* = 4. (**I**) RT-qPCR-based representation of relative changes in the epithelial mRNA of the indicated genes, comparing 1NMPP1 non-treated (“nt”), topical tavilermide (MIM-D3) or THX-B treated to 1NMPP1-injeccted alone or with topical NGF, TrkA^F592A^ mouse corneas. (**J**), (**K**) Split immunofluorescent images of cultured primary hLESCs for TrkA, in (J), and p75^NTR^, in (K), in ΔNp63 that colocalize with Krt-15. Scale bar 50 μm. (**L**) Bright field images of fixed rhodamine stained non-treated or the indicated compounds-treated cultured hLESC cultures. Purple staining intensity indicates size and density of hLESC clones. (**M**) Quantitative representation of colonies forming efficacy (CFA) as per (L). (**N**), (**O**). Normalized RT-qPCR-based relative changes in mRNA of the indicated corneal epithelial cells in (N) and LESCs in (O) marker genes, in primary non treated or treated (as in (**L**)) cultured hLESCs. *n* = 4, ANOVA.

Based on prior evidence suggesting a direct regulatory role of NGF signaling in LESC activity ^17,18,44^, we hypothesized that this regulation involves both TrkA and p75^NTR^ receptors. To test this hypothesis, we first examined the biochemical effects of receptor-specific modulation using topical ligands applied to normal rat corneas *in vivo*: rhNGF; the partial and selective TrkA agonist MIM-D3 (tavilermide) (10 μl of 25 mM), which synergizes with NGF to potentiate TrkA homodimerization without activating p75^NTR^ ^51^, and the selective p75^NTR^ inhibitor THX-B ^52,53^ (10 μl of 5 μg/μl). The optimal topical dosing of the reagents was defined experimentally (see further phenotypic results) and based on the previous reports ^51–53^.

Western blots of corneal epithelial lysates showed that exogenous NGF, which binds both TrkA and p75^NTR^, significantly increased phosphorylation of both TrkA and the p75^NTR^ downstream effector JNK, and expression of p75^NTR^ but, surprisingly, did not increase phosphorylation of TrkA downstream effector AKT ^25^. Tavilermide significantly enhanced TrkA and AKT, as well as notably downregulated JNK phosphorylation. THX-B selectively downregulated p75^NTR^ signaling without affecting TrkA signaling (Figs 2E–H and S1A).

To determine the role of TrkA in LESC identity and maintenance, we employed a previously established loss-of-function mouse model (TrkA^F592A^), enabling selective and reversible TrkA inhibition to avoid the off-target effects of pharmacological TrkA inhibitors. In this model, which we and others used in neuronal contexts ^54,55^, a phenylalanine-to-alanine substitution in the ATP binding pocket renders TrkA selectively sensitive to inhibition by the otherwise kinase-inert compound 1NMPP1 ^55^. Effective TrkA inhibition in corneal epithelial cells was confirmed by western blot analysis of corneal epithelial lysates with anti-pTrkA antibody following systemic 1NMPP1 administration (100 μl of 20 μM). Inhibition of TrkA activity increased the phosphorylation of the p75^NTR^ effector JNK (Fig. S1B,C).

RT-qPCR analysis of TrkA-inhibited corneas revealed significant downregulation of LESC markers (ABCG2, Krt-15, p63), the quiescence marker Gpha2, the epithelial cell marker Krt-12, and the epithelial proliferation marker Ki67 ^3,50^, suggesting that TrkA activity is required for LESC proliferation and differentiation. Topical treatment of TrkA-inhibited corneas with rhNGF, which in this condition only stimulates p75^NTR^ signaling (Fig. S1B,C), restored expression of LESC and quiescence markers but did not rescue downregulation of epithelial and proliferation marker expression (Fig. 2I). Consistently, in TrkA-active cornea, tavilermide upregulated the epithelial cell marker Krt-15, while either tavilermide or THX-B (that enables endogenous NGF to only activate TrkA (Figs 2E-H and S1A)) upregulated expression of the proliferation marker Ki67 (Fig. 2I). These results further suggest that TrkA activity is necessary and sufficient for LESC proliferation and differentiation, while p75^NTR^ activity antagonizes these processes. To assess the relevance of receptor modulation in human cells, primary human (h)LESCs were isolated from cadaver corneas and assessed for expression of TrkA, p75^NTR^, and LESC markers ΔNp63 and Krt-15 (Fig. 2J,K). Clonogenicity assays demonstrated that NGF modestly increased colony number and size, consistent with previous reports ^17^. In contrast, and consistently with the *in vivo* Ki67 expression results (Fig. 2I), THX-B significantly enhanced clonogenicity (the ability of an LESC to divide and form colonies), and tavilermide nearly doubled colony formation, indicating that TrkA activity regulates the proliferation and number of LESCs (Fig. 2L,M). In agreement with the cornea-derived results (Fig. 2I), RT-qPCR analysis further showed that THX-B or tavilermide increased expression of the differentiated epithelial cell marker Krt-12, while TrkA stimulation by NGF or tavilermide significantly upregulated LESC marker genes ABCG2, Krt-15 and p63 (Fig. 2N,O).

Collectively, these results suggest that NGF regulates LESC activity through differential activation of TrkA and p75^NTR^. TrkA signaling promotes LESC proliferation and differentiation, whereas p75^NTR^ antagonizes these effects by suppressing proliferation and differentiation, thereby contributing to maintenance of the stem cell state.

### TrkA and p75^NTR^ oppositely affect corneal epithelial renewal

The above results suggesting that TrkA is required for LESC proliferation and differentiation together with the reported upregulation of local NGF expression following experimental corneal deepithelialization ^12,18^, led us to hypothesize that upregulation of TrkA activity is responsible for accelerated epithelial renewal following corneal wounding ^56^. This hypothesis was further supported by the observation of significant upregulation in TrkA and AKT phosphorylation in response to experimental corneal deepithelialization (Fig. S2A,B). To test this hypothesis directly, we examined whether pharmacological modulation of NGF receptors affects epithelial healing using gain-of-function experiments. Experimentally de-epithelialized rat corneas were topically treated with NGF, tavilermide, or vehicle, and wound closure was monitored starting 12 hours after injury and every 4 hours thereafter. Treatments were administered at each time point. As observed in normally innervated corneas, complete healing occurred within 24 hours in NGF-and vehicle-treated corneas. Selective activation of TrkA with tavilermide accelerated epithelial recovery, resulting in complete wound closure within 20 hours. In all three conditions, wound healing was accompanied by maintenance of corneal transparency (Fig. 3A,B,J).

**Figure 3.**
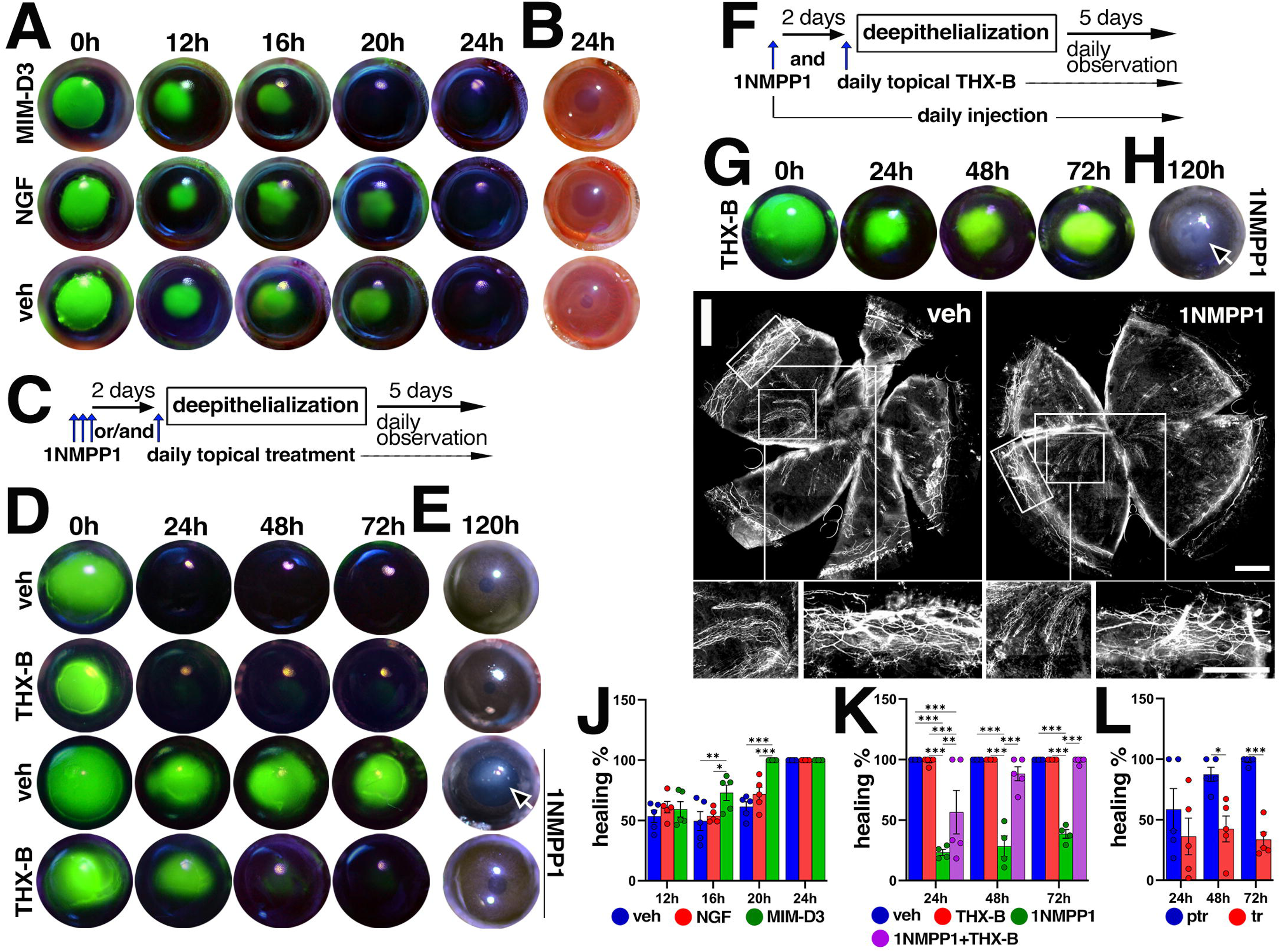
TrkA and p75^NTR^ oppositely affect corneal epithelial renewal. (**A**), (**D**), (**G**) Representative live photos, taken at the indicated time points post de-epithelialization, of de- epithelialized rat in (A) or TrkA^F592A^ mouse in (D) and (G) fluorescein-stained corneas, exposed to the indicated experimental conditions. 0h indicates cornea condition immediately after epithelial removal. The indicated TrkA^F592A^ mice in (D) and (G) were daily intraperitoneally (IP) injected with 1NMPP1, as schematically depicted in (C) and (F), respectively. The corneas were topically treated with the indicated compounds or vehicle alone (“veh”) at every indicated time point in (A), or as per (C or F) in (D or G), respectively. (**B**), (**E**), (**H**) Representative live bright field images at the indicated time points post de-epithelialization, as per (A), (D), (G), respectively. Arrows denote corneal opacification. (**I**) Images of whole mount βIII-tubulin immunolabeled corneas of IP injected with vehicle alone (left panel) or 1NMPP1 (right panels), as per (C), mice, indicate comparable innervation of the central and the limbal areas (magnified framed areas – lower panel). Scale bars in both panels – 500 µm. (**J**), (**K**), (**L**) Quantitative representation of the corneal healing as in (A), (D), (G), respectively. In (L) “ptr” – pretreated, “tr” – treated. n ≥ 5, ANOVA.

To complement these findings, we performed loss-of-function experiments using pharmacogenomic and pharmacological approaches to selectively inhibit TrkA and/or p75^NTR^. TrkA^F592A^ mice were treated with daily intraperitoneal injections (IP) of 1NMPP1 for three subsequent days, including the day of corneal de-epithelialization. Based on the reported kinetics and systemic stability of 1NMPP1 ^55^, this regimen was expected to induce sustained inhibition of TrkA. To inhibit p75^NTR^, corneas of 1NMPP1-pre-treated or untreated TrkA^F592A^ mice were topically administrated with THX-B (Fig. 3C). Corneal healing was monitored daily by fluorescein staining. Complete epithelial recovery occurred within 24 hours in TrkA-active corneas. In contrast, TrkA pre-inhibition abolished epithelial healing and induced severe corneal opacification (Fig. 3D,E,K), comparable to denervated corneas in our model (Fig. 1C,D) and in NK patients ^4,13,14^, suggesting TrkA activity is essential for corneal epithelial renewal.

Strikingly, concomitant treatment of TrkA-pre-inhibited corneas with topical THX-B fully restored epithelial regeneration, resulting in complete healing within 48 hours. As a negative control and to validate TrkA inhibition, topical treatment of 1NMPP1-pre-treated mice with tavilermide failed to restore healing (Fig. S2C-E). The unexpected recovery of TrkA-pre-inhibited corneas may reflect residual activity of the 1NMPP1-treated TrkA^F592A^ hypomorph ^55^, which becomes sufficient to support epithelial repair once p75^NTR^ signaling is suppressed. Alternatively, reversible inhibition of TrkA within TrkA–p75^NTR^ heterodimeric complexes ^57,58^ may be relieved upon p75^NTR^ inactivation and complex dissociation. Indeed, western blot analysis demonstrated that some TrkA phosphorylation was observed in 1NMPP1 treated TrkA^F592A^ corneas, and cotreatment with 1NMPP1 and topical THX-B to inhibit p75^NTR^ enhanced this phosphorylation (Figs. S1B,C and S2F). To test the hypothesis that the observed healing occurred due to residual TrkA activity, phenotypically, we modified the experimental design to ensure sustained TrkA inhibition by administering 1NMPP1 daily throughout the experiment (Fig. 3F). Under these conditions, p75^NTR^ inhibition failed to restore epithelial healing in TrkA-inhibited corneas (Fig. 3G,H,L), confirming the critical requirement for TrkA activity in the epithelial regeneration.

Because TrkA and p75^NTR^ are known regulators of sensory and sympathetic nerve development ^25,37^, we considered whether TrkA inhibition might impair corneal innervation and thereby indirectly induce an NK phenotype. It did not, as immunohistochemical analysis with βIII tubulin revealed comparable limbal and epithelial innervation in TrkA-inhibited and control corneas (Figs 3I and S2G). This finding, together with preserved blink reflex (not shown), indicates that the observed NK phenotype arises from inhibition of non-neuronal TrkA rather than secondary to neurodegeneration.

Collectively, these results demonstrate that (i) TrkA and p75^NTR^ exert opposing effects on corneal epithelial renewal, (ii) TrkA activity is essential for activation of this process, and (iii) corneal innervation alone is insufficient to sustain epithelial renewal, supporting a direct regulatory role of NGF signaling in LESCs.

### Corneal denervation causes upregulation of p75^NTR^ signaling and disrupts LESC homeostasis

Based on the correlation between the TrkA inhibition–induced NK phenotype (Fig. 3D,E) and the regulatory effects of NGF receptor modulation on LESC activity (Fig. 2), we asked whether corneal denervation affects epithelial NGF receptor expression and/or activity. We previously performed scRNA-seq studies comparing de-epithelialized, denervated and control rat corneas ^12^. Analysis of these data indicated that p75^NTR^ expression is upregulated in LESCs following corneal denervation (Fig. 4A and Fig. S3A–D). Consistent with the scRNA-seq results, western blot analysis of mouse corneal epithelial lysates demonstrated that denervation significantly increased p75^NTR^ expression, accompanied by increased phosphorylation of JNK and reduced activation of AKT ^25^ (Fig. 4B,C). Previous studies have reported reduction in LESC number following denervation ^10^. Consistent with this observation, RT-qPCR analysis of corneas four days post-denervation showed significant downregulation of mRNA for the LESC markers (ABCG2, Krt-15, p63), epithelial marker Krt-12, and proliferation marker Ki67 (Fig. 4D). Further analysis of our previous scRNA-seq dataset ^12^ revealed that corneal denervation was associated with reduced LESC and epithelial cell populations, and an increase in TAC and mesenchymal cell populations (Fig. 4E).

**Figure 4.**
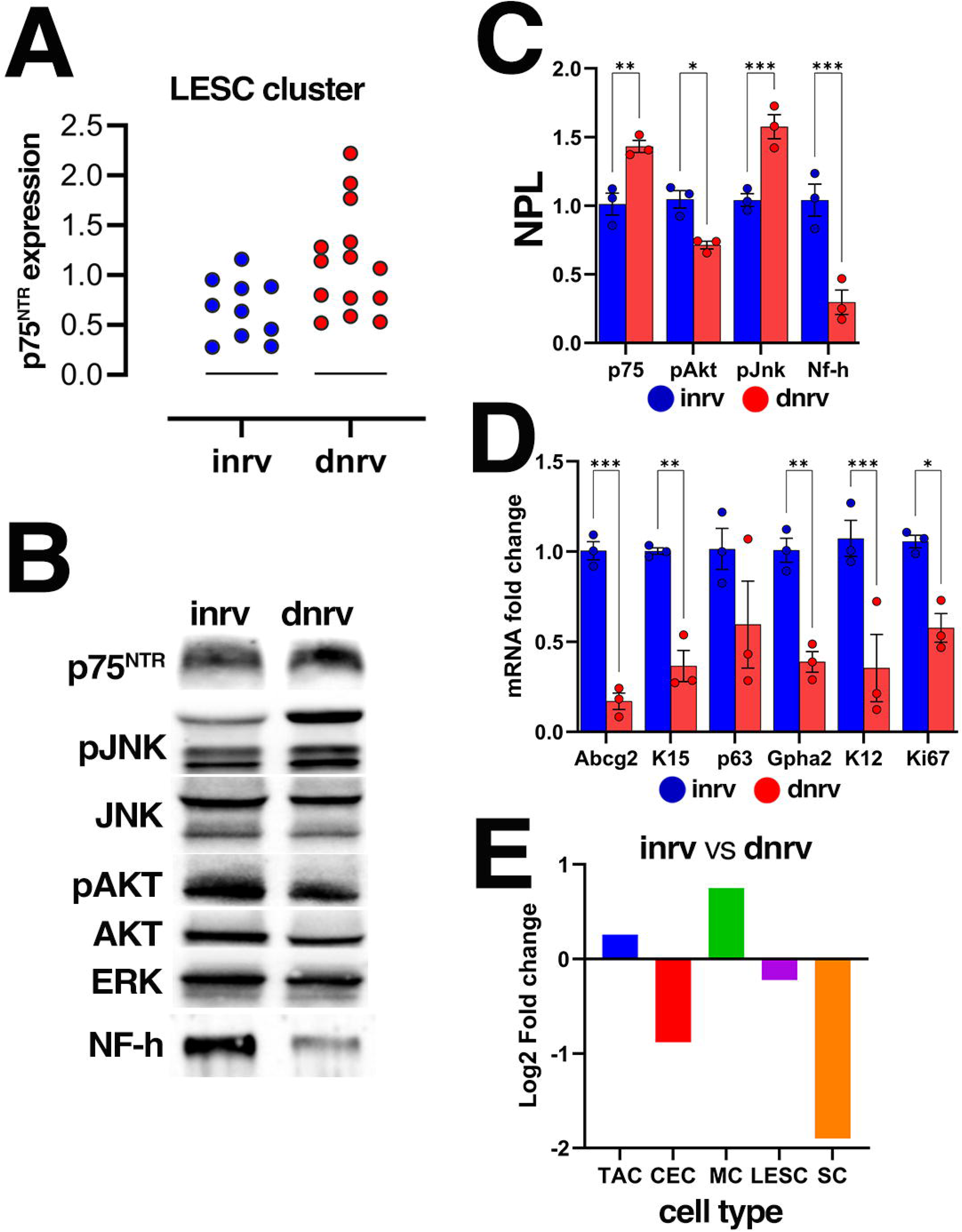
Corneal denervation causes upregulation of the inhibitory p75^NTR^ axis. **(A**) scRNAseq-based ^12^ violin plot presentation of *p75^NTR^*mRNA expression in innervated (“inrv”) and five days post-denervated (“dnrv”) rat corneas. (**B**) Western blot analysis of innervated (inrv) or five days post-denervated (dnrv) rat corneal epithelial lysates for the indicated proteins and their phosphorylated form (p). Heavy chain neurofilament “NH-h” indicates significant denervation. (**C**) Quantitative representation of normalized protein level (NPL) as per (B). (**D**) RT-qPCR-based representation of relative changes in the epithelial mRNA of the indicated genes following denervation. [(C), (D)] *n* = 3, T-test. (**E**) scRNAseq-based ^12^ representation of the corneal denervation-caused change in the volume of the transient amplifying calls (TAC), corneal epithelial cells (CEC), mesenchymal cells (MC), limbal epithelial stem cells (LESC) and Schwann cells (SC).

Collectively, these findings indicate that corneal denervation shifts homeostatic NGF signaling toward p75^NTR^ dominance and is associated with suppression of LESC maintenance and activity, as seen with TrkA inhibition (Fig. 2I).

### Selective TrkA activation rescues epithelial renewal in NK corneas

Based on our results that denervation shifts the balance of TrkA/p75^NTR^ signaling towards p75^NTR^ in LESCs, we hypothesized that in denervated corneas, pharmacological modulation of NGF receptors in favor of the TrkA axis would restore epithelial regenerative capacity. To test this hypothesis, we assessed the effect of topical treatment of denervated and de-epithelialized rat corneas, as in Fig 1A, either by inhibiting p75^NTR^ by topical THX-B or activating TrkA with tavilermide.

Supporting this hypothesis, both treatments induced complete epithelial healing of otherwise non-healing denervated corneas within 24 hours, recapitulating the healing observed in normally innervated corneas (Fig. 5A,B,I). Note that treatment with THX-B alone did not affect healing of non-denervated control corneas (Fig. S4A,B). As observed with rhNGF treatment (Fig. 1B), βIII tubulin immunostaining of treated corneas showed no evidence of reinnervation (Fig. S4C), indicating that epithelial recovery was driven by direct effects on epithelial cells, independent of corneal nerves.

**Figure 5.**
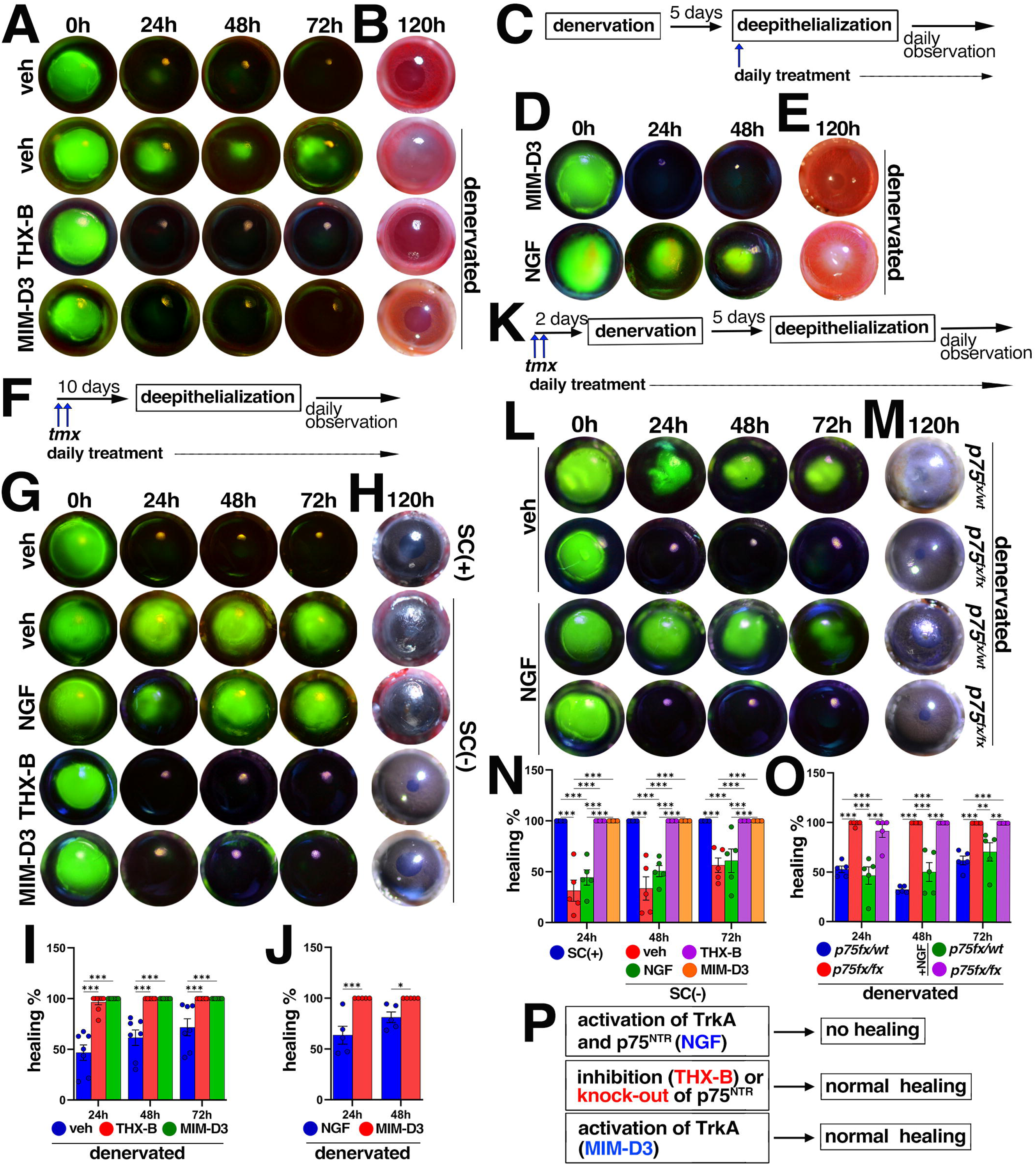
**Domination of the TrkA axis induces epithelial renewal of NK cornea**. (**A**), (**D**), (**G**), (**L**) Representative live photographs of de-epithelialized fluorescein-stained normally innervated or denervated rat in (A) (as per schematic description in Figure 1A) and (D), or Schwann cell present (*Sox10-TdT*) or ablated (*Sox10-TdT-DTA*) in (G), or denervated p75^NTR^ heterozygous (*p75^fx/wt^*) or null (*p75^fx/fx^*) in (L) mouse corneas, demonstrating the course of the epithelial healing in response to topical treatment with the indicated compounds or vehicle alone (“veh”). 0h indicates corneal condition five days post denervation (immediately after de-epithelialization). (**B**), (**E**), (**H**), (**M**) Representative live bright field images of the corneas as per (A), (D), (G), (L), respectively, 120 hours post-deepithelialization. (**C**), (**F**), (**K**) Schematic description of the experiments design in (D), (G), (L), respectively. (**I**), (**J**), (**N**), (**O**) Quantitative representation of corneal healing as per (A), (D), (G), (L), respectively. Each data point represents the fluorescein-negative (healed) area. *n* ≥ 5, ANOVA. (**P**) Conceptual summary of the effect of the NGF receptors modulation (as per (A), (L) and Figure 1C) on healing of NK cornea.

The results of treatment initiated at the time of corneal denervation (Figs. 1A and 5A,B), although demonstrating proof-of-concept for restoring epithelial healing in anesthetic corneas, do not address the more clinically-relevant chronic NK. We therefore repeated the experiment using a delayed treatment paradigm, in which tavilermide was administered at the dose applied in clinical trials (50 μg/μl × 10 μl) ^59^ following epithelial injury five days after denervation (Fig. 5C), when the NK phenotype is already established (^12^ and in Fig. 5A,B). Under these conditions, tavilermide induced rapid and complete epithelial healing. In contrast, topical NGF administered at the clinically daily dose (0.5 μg/μl in 10 μl) failed to promote healing (Fig. 5D,E,J). These findings support selective TrkA activation as a potential therapeutic strategy for NK.

We previously demonstrated that Schwann cells associated with corneal sensory nerves mediate innervation-dependent epithelial renewal ^12^. To further investigate their role in NGF signaling, we examined the effect of receptor modulation in Schwann cell–deficient corneas. Using a *Sox10-iCre^ERT^*^2^*^/+^;R26-LSL-TdT;R26-LSL-DTA* (*Sox10-TdT;DTA*) mouse model, in which topical tamoxifen induces expression of diphtheria toxin resulting in targeted Schwann cell ablation, we induced local Schwann cell loss with three consecutive daily topical tamoxifen treatments ^12^. Corneas were subsequently treated with THX-B, rhNGF, tavilermide, or vehicle. Ten days after induction, corneas were de-epithelialized and healing was assessed (Fig. 5F).

As previously reported ^12^, near-complete ablation of TdTomato-positive Schwann cells elicited the NK phenotype in vehicle-treated *Sox10-TdT;DTA* mice (Fig. 5G,H; Fig. S5A). Consistent with denervation experiments, rhNGF failed to rescue epithelial healing. In contrast, THX-B or tavilermide fully restored epithelial regeneration, comparable to de-epithelialized control corneas in which Schwann cells were present (Fig. 5G,H,N).

Although these findings indicate robust rescue of epithelial healing, they do not directly demonstrate LESC-specific effects. Besides LESCs, p75^NTR^ is expressed by sensory neurons and the associated Schwann and mesenchymal cells ^37,60,61^ (Fig. S3C). Therefore, to focus on LESCs, we developed a conditional p75^NTR^ knockout LESC mouse model. A Krt14-Cre construct was previously used for induced LESC specific gene expression in mice ^62^. To selectively ablate p75^NTR^ in LESCs, we crossed *Krt14-Cre^ERT/+^* with *p75^NTR-FX^* ^63^ to generate *Krt14-Cre^ERT/+^;p75^NTR-FX^* (*K14-p75^fx/fx^*) mice. Tamoxifen-induced p75^NTR^ deletion was validated by immunohistochemistry and RT-qPCR (Fig. S5B-D).

LESC-specific p75^NTR^ deletion did not affect corneal homeostasis under baseline conditions (not shown). However, in denervated (Fig. S5E) and de-epithelialized corneas, homozygous p75^NTR^ deletion resulted in complete epithelial recovery within 24 hours, whereas heterozygous controls failed to heal. Notably, normal healing was observed in the p75^NTR^ null rhNGF treated corneas (Fig. 5L,M,O), demonstrating that p75^NTR^ mediates the inhibitory effect of NGF in LESCs. Collectively, these results demonstrate that, unlike rhNGF, selective pharmacological activation of TrkA or inhibition, or LESC-specific ablation of p75^NTR^ restores epithelial renewal in NK corneas (Fig. 5P), highlighting the central role of NGF receptor balance in regulating LESC activity. Furthermore, these findings validate the previously identified role of Schwann cells ^12^ in mediating innervation-dependent corneal epithelial renewal.

### The p75^NTR^-induced NK phenotype is ligand dependent

It was previously reported that topical NGF neutralizing antibody (αNGF) delays healing of experimentally de-epithelialized rat cornea, speculated to be due to NGF blockade–induced inhibition of TrkA ^18^. However, since αNGF also interferes with binding of both NGF and proNGF to p75^NTR^ ^64^, and considering the results above suggesting involvement of both NGF receptors in regulation of LESCs, we realized that the previously proposed model required rethinking.

Selective activation of TrkA or inhibition of p75^NTR^ supports LESC proliferation and clonogenicity to a greater extent than exogenous NGF (Fig. 2I,L,M). Therefore, together with previously suggested autocrine NGF signaling in LESCs ^17^, we hypothesized that either NGF blockade or exogenous activation of p75^NTR^ would interfere with LESC proliferation. Supporting this hypothesis, treatment with either αNGF or recombinant proNGF almost completely blocked colony formation by cultured primary hLESCs (Fig. 6A,B). To validate this effect *in vivo*, we treated deepithelialized, normally innervated rat corneas with one or two daily doses of αNGF (10 μl of 50 ng/ml) or recombinant proNGF (10 μl of 50 µg/ml). In agreement with the *in vitro* experiment, αNGF delayed epithelial healing in a dose-dependent manner with complete healing observed at 48 or 72 hours following one or two daily doses, respectively, and induced corneal opacification, likely reflecting the extent of partial blockade of endogenous NGF. In contrast, proNGF completely prevented corneal healing and induced NK-like severe opacification (Fig. 6C-F and Fig. S6A,B,D). These findings support previously described autocrine NGF signaling in LESCs ^17,49^ and indicate that p75^NTR^-mediated suppression of proliferation is ligand-dependent.

**Figure 6.**
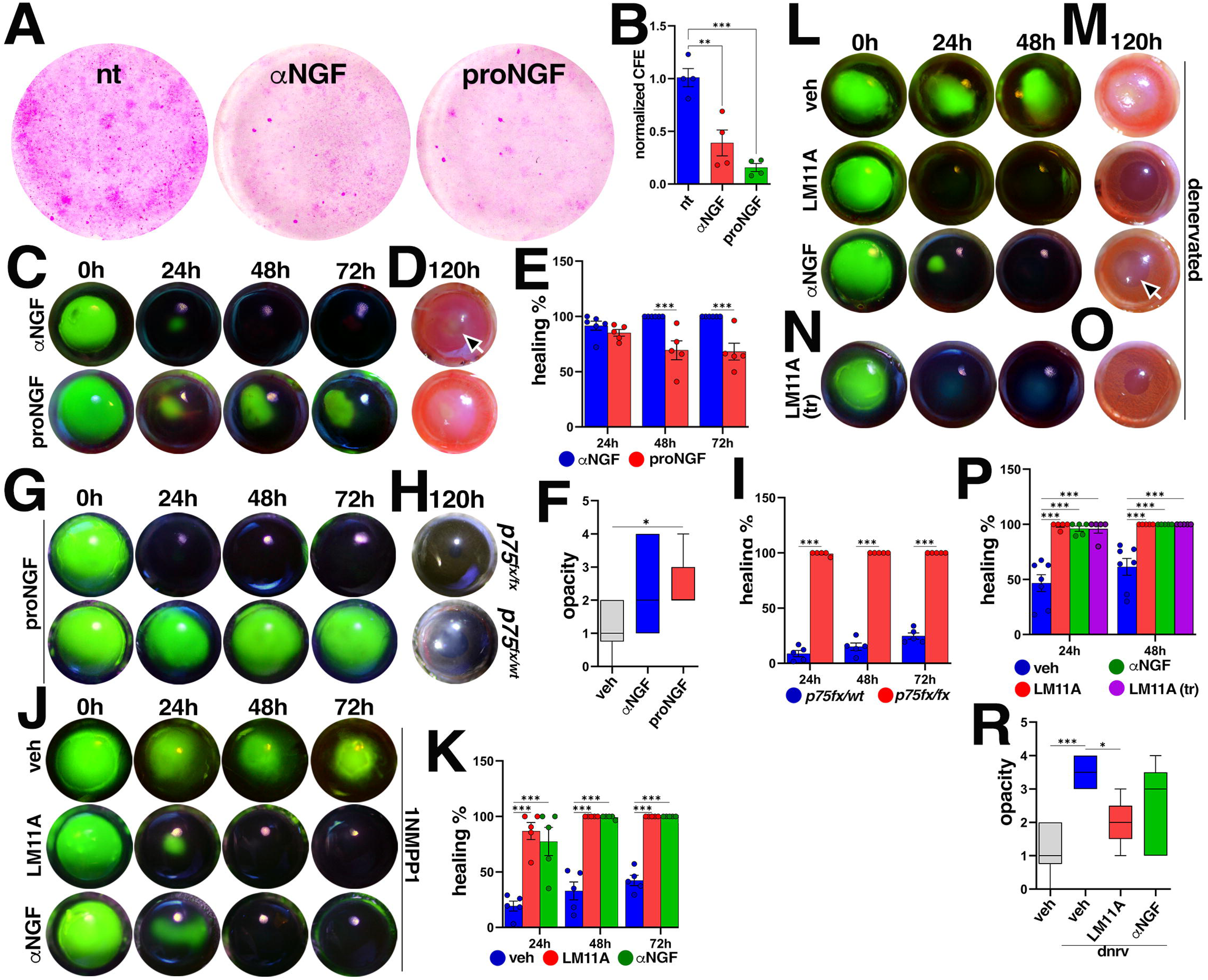
The p75^NTR^-induced NK phenotype is ligand dependent. (**A**) Bright field images of fixed rhodamine stained non-treated (“nt”) or the indicated compounds-treated cultured hLESC cultures. Purple staining intensity indicates size and density of hLESC clones. (**B**) Quantitative representation of CFA stands for colonies forming efficacy in (A). *n* = 4. (**C**), (**G**), (**J**), (**L**), (**N**) Representative live photographs of fluorescein-stained de-epithelialized innervated in (C) or denervated in (L) and (N) rat, or innervated p75^NTR^ heterozygous (*p75^fx/wt^*) or null (*p75^fx/fx^*) topical tamoxifen pre-treated in (**G)**, 1NMPP1 intraperitoneally pre-treated TrkA^F592A^ (as per Figure 3C) in (J) mouse corneas, demonstrating the course of the epithelial healing in response to topical treatment with the indicated compounds or vehicle alone (“veh”). The daily treatment started at the day of corneal deepithelialization for (C), (G), (J), (N) or the day of corneal denervation for (L). The experimental design for (L) and (M), as per the schematic description in Figures 1A and 5C, respectively. (**D**), (**H**), (**M**), (**O**) Representative live bright field images of the corneas as per (C), (G), (L), (N), respectively, 120 hours post-de-epithelialization. Arrows in (D) and (M) denote opacified healed corneas. (**E**), (**I**), (**K**), (**P**) Quantitative representation of corneal healing as per (C), (G), (J) or (L and N), respectively. Each data point represents the fluorescein-negative (healed) corneal area. In (P) “tr” indicates for treatment with LM11A-31 started on the day of deepithelialization, five days after denervation in (N). (**F**), (**R**) A deep learning architecture pretrained convolutional neural network (CNN) analysis-based quantitative representation of the corneal opacity as in (C), (L), respectively, comparing between the experimental conditions and vehicle alone treated normally innervated de-epithelialized (“dnrv”) corneas (as in Figures 1C or 5A). [(E-I), (K), (P), (R)] *n* ≥ 5, ANOVA.

To further validate this mechanism in a loss-of-function assay, we assessed the effect of proNGF on the corneal healing of LESC-specific p75^NTR^-deficient mice. Similar to the rat experiments (Fig. 6C,D), proNGF treatment prevented corneal healing in tamoxifen-induced heterozygous K14-p75^fx/wt^ mice. In contrast, proNGF had no effect on epithelial healing in LESC-specific p75^NTR^-null corneas (Fig. 6G-I), providing complementary evidence for ligand-dependent inhibitory activity of p75^NTR^.

Above we showed that inhibition of p75^NTR^ restored healing in TrkA-pre-inhibited corneas (Fig. 3C-E,K). Therefore, we hypothesized that blocking NGF/proNGF binding to p75^NTR^ would similarly rescue epithelial regeneration under TrkA pre-inhibition. A single daily dose of topical αNGF was compared to the selective p75^NTR^ antagonist LM11A-31 ^65,66^. TrkA inhibition was induced by systemic 1NMPP1 treatment of TrkA^F592A^ mice prior to corneal de-epithelialization (Fig. 3C). Consistent with THX-B treatment (Fig. 3D,E), both LM11A-31 (10 μl of 25µg/μl) and αNGF restored epithelial healing of TrkA-pre-inhibited corneas within 48 hours. In the same experimental paradigm, αproNGF (10 μl of 0.7 μg/ml), which specifically blocks proNGF binding to p75^NTR^ ^64^, induced near complete healing with delayed kinetics of 72 hours (Fig. S6C,E). These results demonstrate that p75^NTR^-mediated inhibition of epithelial renewal is dependent on endogenous ligand availability and suggest that both mature NGF and proNGF contribute to receptor activation.

Previous studies, including our own ^10,12^, demonstrated that corneal denervation induces upregulation of local NGF expression (Fig. S6F-H). Based on this, we hypothesized that p75^NTR^ dominance in denervated corneas is driven by increased endogenous NGF. To test this, we examined the effect of topical LM11A-31 p75^NTR^ antagonist or αNGF in denervated, de-epithelialized rat corneas. The rationale for using αNGF was the observed p75^NTR^ dominance following denervation (Fig. 4A-C), along with the epithelial healing in αNGF-treated innervated rat and TrkA pre-inhibited mouse corneas (Fig. 6C-E,J,K) that is indicative of a dose dependent, partial block of NGF. However, unlike LM11A-31, which induced healing within 24 hours, αNGF-mediated healing was delayed by 24 hours and associated with corneal opacification, likely due to partial inhibition of TrkA pro-regenerative signaling along with p75^NTR^ degenerative signaling (Figs 6L,M,P,R and S6I). Moreover, similar and consistent with tavilermide, LM11A-31 induced epithelial healing when applied in the chronic model (five days post-denervation) (Figs 5C and 6N,O), demonstrating its clinical relevance. As expected, LM11A-31 did not affect healing of normally innervated de-epithelialized corneas (not shown). Collectively, these findings demonstrate that p75^NTR^-mediated inhibition of epithelial renewal in both innervated and denervated corneas is NGF-dependent. Importantly, these results suggest that the NK phenotype arises not from NGF deficiency but from excess endogenous NGF–driven p75^NTR^ activation. Furthermore, they identify p75^NTR^ inhibition as a rational therapeutic strategy for NK.

## DISCUSSION

### Opposing NGF-mediated TrkA and p75 ^NTR^ signaling regulates LESC activity

Homeostatic corneal epithelial renewal is normally slow and sustained by long-term LESC self-renewal, whereas epithelial injury triggers rapid re-epithelialization through accelerated LESC activation and migration of newly generated epithelial cells ^8,44^. The presence of two NGF receptors in LESCs (Fig. 2A–D,J,K) with distinct and partially antagonistic signaling properties and binding kinetics ^67,68^, which can act independently or cooperatively depending on their environment, provides a plausible mechanism for this dynamic regulation.

We demonstrate that selective activation of TrkA with the agonist tavilermide, similar to NGF, increased expression of LESC-associated markers and stimulated hLESC proliferation more strongly than NGF (Fig. 2I,L-O). Conversely, TrkA inhibition *in vivo* reduced expression of LESC maintenance, differentiation, and proliferation markers (Fig. 2I), whereas NGF neutralization with αNGF ^17,54^ suppressed hLESC proliferation and delayed healing, and caused opacification of experimentally de-epithelialized corneas in a dose-dependent manner (Figs. 6A-E and S6A,B,D).

Consistent with previous reports of injury-induced NGF upregulation in multiple tissues including the cornea ^18^, we further observed increased NGF expression by Schwann cells following corneal de-epithelialization in rats ^12^. Given the high sensitivity of the TrkA signaling-driven NGF receptor complex to availability and changes in NGF ^67–69^, and consistent with the observed upregulation of TrkA phosphorylation in response to experimental deepithelialization (Fig. S2A,B), these findings suggest that TrkA is involved in regulating the extent and the rate of proliferation of LESCs in response to injury. Validating these hypotheses, in contrast to exogenous NGF, tavilermide, which increases TrkA sensitivity to NGF and stabilizes the preformed homodimers ^51^, significantly accelerated recovery of experimentally de-epithelialized corneas (Fig. 3A,B). Consistently, epithelial healing was blocked by either TrkA inhibition or p75^NTR^ activation, and restored by p75^NTR^ inhibition in TrkA-pre-inhibited cornea (Figs. 3C-E and 6C-K). These further suggest TrkA as a stimulator of LESC activity, defining the extent and the rate of their proliferation and differentiation in response to changes in the epithelial integrity. Here we demonstrate that activation of p75^NTR^ with NGF reversed LESC marker expression in TrkA-inhibited corneas without restoring epithelial differentiation and proliferation marker expression (Fig. 2I), while selective activation of p75^NTR^ with proNGF markedly inhibited hLESC proliferation (Fig. 6A,B). Consistently, pharmacological inhibition of p75^NTR^ enhanced proliferation and differentiation marker expression *in vivo* and *in vitro* (Fig. 2I,L-O). Notably, competitive inhibition of p75^NTR^ with LM11A-31 (Fig. 6J,K), comparable to THX-B (Fig 3C-E,K), promoted epithelial healing in TrkA-pre-inhibited de-epithelialized corneas. Likewise, daily topical administration of αNGF or αproNGF albeit with delayed kinetics induced epithelial healing in TrkA-pre-inhibited corneas *in vivo* (Figs 6J,K and S6C,E). These observations identify p75^NTR^ as an inhibitor of LESC proliferation and differentiation, whose activity is dependent on endogenous NGF.

Collectively, our findings support a model in which NGF controls corneal epithelial renewal through activation of TrkA-p75^NTR^ receptor complex in LESCs, with TrkA supporting LESC proliferation, differentiation, and probably migration ^44^, whereas p75^NTR^ restrains these processes to maintain stem cell homeostasis. Following corneal injury, the rapid upregulation of local NGF further shifts receptor occupancy toward TrkA, which we suggest amplifies AKT-driven proliferation and migration, thereby promoting epithelial regeneration and restoration of the corneal surface.

### Corneal innervation regulates epithelial renewal through NGF-dependent TrkA–p75^NTR^ signaling

Beyond their protective role, the Schwann cell-insheathed corneal nerves are an integral component of the limbal niche and contribute to both homeostatic and injury-induced epithelial renewal ^4,12^. Impaired healing observed in experimentally denervated corneas is associated with LESC loss ^10^ (Fig. 4E), underscoring the critical role of the nerves in maintaining LESCs. The findings presented here, together with the previously generated scRNAseq datasets ^12^, establish a direct mechanistic link between corneal innervation and NGF-regulated epithelial renewal. We show that NGF’s effects on LESCs are not intrinsically pro- or anti-regenerative but rather are context-dependent and defined by the relative balance of TrkA and p75^NTR^ activity, a balance that is maintained by corneal innervation and inverted by denervation.

Unexpectedly, and in contrast to the widely accepted pro-regenerative effect of NGF ^20,45^, topical rhNGF failed to promote healing of completely denervated or Schwann cell ablated and de-epithelialized corneas. In contrast to exogenous NGF, yet agreement with the demonstrated critical role of TrkA (Fig. 3D,E,K), selective stimulation of the TrkA with tavilermide, prevented NK development and promoted rapid epithelial healing (Figs 1C,D and 5).

What accounts for these opposing phenotypic outcomes? Our data show that corneal denervation induced p75^NTR^ mRNA upregulation in LESCs (Fig. 4A), correlating with increased p75^NTR^ expression and downstream signaling, and reduced TrkA signaling in rat corneal epithelium (Fig. 4B,C). Consistent with TrkA inhibition (Fig. 2I), in our *in vivo* experimental system, corneal denervation suppressed expression of LESC maintenance, differentiation, and proliferation markers (Fig. 4D) and correlated with reduced LESC and epithelial cell volume (Fig. 4E). These findings suggest that denervation shifts the TrkA–p75^NTR^ signaling balance toward p75^NTR^, thereby contributing to the failure of epithelial renewal.

Correlating with the p75^NTR^ dominance and ligand dependent activity, we and others observed a marked increase in local NGF expression following corneal denervation ^10,12^ (Fig. S6F-H). Here we demonstrate that either non-competitive or competitive inhibition of p75^NTR^ with THX-B or LM11A-31, respectively (Figs 5A,B,I and 6L,M,P), or its genetic ablation in LESCs (Fig. 5L,M,O) promoted epithelial healing of denervated de-epithelialized corneas. These findings demonstrate that in denervated cornea endogenous NGF-dependent p75^NTR^ activity is responsible for the failure of epithelial healing, characteristic of NK.

Consistently, daily topical administration of αNGF, albeit with delayed kinetics, induced the epithelial healing of denervated corneas (Fig. 6L,M,P). While αNGF interferes with activation of both receptors, its rescuing effect is consistent with the distinct ligand-binding kinetics of TrkA and p75^NTR^, whereby p75^NTR^ exhibits rapid ligand association–dissociation and TrkA displays slower binding dynamics ^67^, rendering p75^NTR^ more sensitive to fluctuations in NGF availability. At the same time, the less specific endogenous TrkA agonist NT3 might also contribute to the retained TrkA activity (as shown for innervated αNGF-treated cornea Fig. S6I) in the condition of NGF neutralization-mediated uncoupling of the TrkA-p75^NTR^ complex ^34,39^. Although, this sub-optimal TrkA activity likely contributes to the delayed corneal healing and the epithelial, and stromal opacification (Fig. 6L,M,P,R). The latter is indicative of involvement of NGF signaling in supporting stromal clarity independently of its role in epithelial regeneration, although this mechanism was not directly addressed in the present study.

Collectively, our results establish a direct mechanistic link between corneal innervation and epithelial renewal and suggest a conceptual model: in the innervated cornea, where endogenous NGF is present at physiological concentrations, TrkA-p75^NTR^ receptor complexes generate high-affinity NGF binding sites that favor sustained TrkA-AKT signaling ^36,70–72^. In contrast, by increasing the proportion of rapidly cycling p75^NTR^, denervation shifts the balance of NGF toward p75^NTR^-JNK, thereby causing NK phenotype. The two-receptor system therefore functions not merely as an on-off switch but as a dynamic integrator of neural and epithelial signals, capable of calibrating LESC activity in response to both injury and innervation status. Our findings further suggest a distinct Schwann cell–dependent mechanism regulating p75^NTR^ expression in LESCs. This hypothesis is being addressed in our ongoing research.

### Rebalancing TrkA and p75^NTR^ as a therapeutic strategy

The pivotal REPARO trial reported corneal healing in 74% of cenegermin-treated patients versus 43% in controls, yet many responses were incomplete and approximately 25% were associated with adverse events including epithelial breakdown and blurred vision ^47^. More recent real-world analyses report low-certainty improvement in re-epithelialization without significant gains in visual acuity ^73^, along with up to 40% recurrence or worsening within four months of treatment ^74–76^. These data collectively highlight the limited and often transient efficacy of cenegermin supplementation in severe NK and underscore the need for a deeper mechanistic understanding of why topical NGF is insufficient in severe NK.

The original rationale for cenegermin use in NK was that exogenous NGF administered at 20 μg/ml, six times daily ^41,43,77^ would restore the local trophic milieu and induce epithelial healing. Arguing against this rationale, we and others showed that corneal denervation does not reduce, but instead increases, local NGF expression ^10,12^ (Fig. S6F-H) that, in turn, amplifies p75^NTR^ activity in LESCs (Fig. 4A-C). Accordingly, and consistent with severe NK case ^42,43,47,78–81^, rhNGF, even at dose of 500 μg/ml (which is 25-fold the clinical concentration), failed to restore epithelial healing in denervated rodent corneas (Fig. 5C-E,J). Moreover, our *in vivo* (not shown) and clinical observations suggest that the supraphysiological rhNGF dosing used in cenegermin NK therapy induces ocular inflammation and neovascularization ^19,47,82^, likely contributing to stromal scarring. In contrast, restoring TrkA dominance with tavilermide, or indirectly via p75^NTR^ inhibition/activation blocking with THX-B or LM11A-31 (Figs 5A,B,J and 6L,M,P), induced rapid and complete epithelial healing, without inducing opacification in the established NK condition *in vivo* (Figs 5C-E,J and 6N-P). Moreover, topical tavilermide and LM11A-31 not only prevented development of the NK phenotype but also restored epithelial healing in de-epithelialized corneas with established NK (Figs Fig. 5D,E,J and 6N-P). Collectively, findings identify a mechanistically distinct therapeutic strategy that targets the receptor-level mechanism rather than supplementing its ligand, with potentially safer therapeutic profile than rhNGF.

Cenegermin frequently induces severe ocular pain ^23^, likely mediated by p75^NTR^ activation ^83,84^. In contrast, selective TrkA activator tavilermide demonstrated established ocular tolerability without significant nociceptive adverse effects in Phase II and III dry eye trials ^59^. LM11A-31’s specific competitive inhibition of p75^NTR^ that avoids TrkA overactivation and its associated pro-inflammatory effects, demonstrated favorable safety profile in Phase IIa trials for Alzheimer’s disease ^66^. Together with the pro-regenerative activity, these properties support the rapid clinical translation of both compounds for NK. Furthermore, given the reported neurotrophic activity of tavilermide ^85^ and the neuroprotective effects of LM11A-31 ^66,86^, both compounds may additionally contribute to restoration of corneal innervation, an effect that if confirmed would further contribute to their therapeutic potential. Whether selective TrkA activation and p75^NTR^ inhibition offer complementary therapeutic benefits, and whether combination approaches provide more durable receptor rebalancing than either monotherapy alone, are important questions for future investigation.

### Summary

Research on corneal epithelial renewal has largely progressed along two independent paths: regulation of LESC and their niche ^16,87–89^ and the role of sensory innervation in epithelial homeostasis and repair ^10,48,90^. Here, we unify these fields by identifying an NGF-dependent signaling mechanism linking corneal innervation to LESC-driven epithelial renewal. We show that nerve-associated Schwann cells maintain epithelial homeostasis by sustaining TrkA dominance over p75^NTR^ signaling in LESCs, whereas denervation shifts this balance toward anti- regenerative p75^NTR^ activity through simultaneous p75^NTR^ upregulation and increased local NGF expression. Following this, we identify selective TrkA activation with tavilermide or p75^NTR^ inhibition with LM11A-31 as mechanistically distinct therapeutic strategies for fully or severely denervated NK corneas.

## MATERIALS AND METHODS

### Study design

This study was designed to define how corneal sensory innervation regulates LESC-mediated epithelial renewal and to determine the mechanistic basis of NGF signaling in neurotrophic keratopathy (NK). We combined *in vivo*, *ex vivo*, and *in vitro* approaches to test whether corneal nerves control epithelial repair through receptor-specific NGF signaling. In a rodent model of complete corneal denervation induced by electrocautery of the ophthalmomaxillary branch of the trigeminal nerve ^48^, we assessed the effects of topical rhNGF, or selective pharmacological TrkA activation or p75^NTR^ inhibition on the epithelial healing. To define receptor-specific mechanisms, we examined TrkA and p75^NTR^ expression and signaling in rodent cornea and human corneal epithelial cells/LESCs using immunohistochemistry, western blotting, RT-qPCR, and scRNA-seq analysis. We further used TrkA^F592A^ loss-of-function mice and LESC-specific p75^NTR^ conditional knockout mice ^12,54,63^ to test receptor function in epithelial renewal. Primary human LESC cultures were used to evaluate clonogenicity and lineage-associated gene expression. Finally, to assess translational relevance, we tested receptor-selective compounds in denervated, Schwann cell–deficient ^12^, and chronically denervated corneas.

### Statical analysis

All statistical analyses were performed using GraphPad Prism software (GraphPad Software, San Diego, CA, USA). For *in vivo* assays animals were randomized to experimental groups where appropriate, and investigators were blinded during qualitative and quantitative analyses. Group sizes of at least five animals per condition were considered sufficient to ensure representative biological reproducibility. No animals or biological samples were excluded from analysis unless otherwise specified. Data are presented as mean ± standard error of the mean (SEM). Comparisons between two groups were performed using unpaired Student’s t tests. Comparisons among multiple groups were performed using one-way analysis of variance (ANOVA) followed by Tukey’s post hoc test, as appropriate. A p-value < 0.05 was considered statistically significant and indicated as follows: \**p* < 0.05; \*\**p* < 0.01; \*\*\**p* < 0.005.

### List of Supplementary Materials

## Materials and Methods

Fig. S1 – S6

## Acknowledgments

We thank Dr Uri Saragovi (McGill University) for consulting on TrkA and p75^NTR^ small molecule modulators and kindly sharing MIM-D3 and THX-B for preliminary observations (later purchased from commercial resources) and proNGF neutralizing antibody that is commercially non-available. We thank Dr Brian Pierchala (Indiana University) for kindly sharing the *p75^NTR-FX^* mouse model. We thank Dr Karen Meerovitch (Mimetogen Pharmaceuticals, QC Canada) for contribution to the manuscript’s revision.

## AUTHORS CONTRIBUTION

K.F., G.H.B., T.G. and A.A. conceived the research.

K.F. conceptualized and designed the study and wrote the original draft.

K.F., G.H.B., R.S.F. and A.A. developed methodology.

K.F. and G.H.B. supervised the research.

A.H., K.T., R.K., K.M., L.C., A.I.Q, X.W. and N.B. performed the experimental work.

D.J.K. developed and performed the AI-based corneal opacification analysis.

K.F., G.H.B., A.H., R.K., J.S.P., T.G., and D.R.K. reviewed & edited the manuscript.

## INTELECTUAL PROPERTY

UNITED STATES PATENT AND TRADEMARK OFFICE.

Utility – U.S. National Stage under 35 USC 317. APPLICATION # 19/581,675, ATTORNEY DOCKET #144578.00493. Title of Invention: Compositions Comprising p75NTR Inhibitors and Recombinant Nerve Growth Factor and Methods for the Treatment of Corneal Diseases and Disorders Using the Same. 04/24/2026 03:01:23 PM Z ET

UNITED STATES PATENT AND TRADEMARK OFFICE.

Utility – Provisional Application under 35 USC 111(b). APPLICATION #63/900,993, ATTORNEY DOCKET #144578.00473. Title of Invention: Methods for Treatment Neurotrophic Keratopathy Using p75 Activation Inhibitors. 10/17/2025 02:50:28 PM ET

UNITED STATES PATENT AND TRADEMARK OFFICE

International Application (PCT) for filing in the US receiving office. APPLICATION # PCT/US25/47085, ATTORNEY DOCKET #144578.00419. Title of Invention: Methods for treating Neurotrophic Keratitis. 09/19/2025 11:50:45 AM Z ET

UNITED STATES PATENT AND TRADEMARK OFFICE

Utility – Provisional Application under 35 USC 111(b). APPLICATION #63/696,424, ATTORNEY DOCKET #144578.00419. Title of Invention: Methods for treating Neurotrophic Keratitis. 09/19/2024 11:07:22 Z ET

UNITED STATES PATENT AND TRADEMARK OFFICE

Utility – Provisional Application under 35 USC 111(b). APPLICATION #63/593,283, ATTORNEY DOCKET #144578.00362. Title of Invention: Compositions Comprising a New Small Molecule and Cenegermin for the Treatment of Corneal Diseases and Disorders Using the Same. 10/26/2023 11:38:32 AM ET

## DATA AND MATERIAL AVAILABILITY

All data associated with this study are present in the paper or Supplementary Materials. The scRNA-seq datasets has been published in Mirmoeini et al 2023. The R scripts used for scRNA-seq is available at <u>Zenodo (</u>https://zenodo.org/records/17638176). All materials used or generated in this study are commercially available or will be supplied upon reasonable request.

