## Supplementary Material for "A Nerve-Dependent NGF Receptor Switch Controls Corneal Epithelial Renewal In Neurotrophic Keratopathy"

**MATERIALS AND METODS**

**Drugs and reagents**

The following pharmacologic agents and reagents were used for *in vivo* and *in vitro* assays: recombinant human β-NGF (rhNGF) (Invitrogen, Cat# A42578), NGF/proNGF neutralizing antibody (αNGF) (Alomone Labs, Cat# ALM-006), recombinant human pro-NGF (Alomone Labs, Cat# N-280), TrkA agonist Tavilermide (MIM-D3, provided by Mimetogen, NY, USA), the hippomorphic TrkA^F592A^ inhibitor 1NMPP1 (MedChemExpress, Cat# HY-13942), p75^NTR^ antagonist LM11A-31 (MedChemExpress, Cat# HY-1117088), p75^NTR^ inhibitor THX-B (MedChemExpress, Cat# HY-137322), proNGF neutralizing antibody (αproNGF; generously shared by Uri Saragovi PhD McGill University). Isoflurane anesthesia (Primal Critical Care, PA, USA) was used for all surgical and imaging procedures. Meloxicam in extended-release polymer (Wedgewood Pharmacy, NJ, USA) was administered for perioperative analgesia. Powder formulations of rhNGF, pro-NGF, αNGF, αproNGF, THX-B or 1NMPP1, or stock tavilermide solution were dissolved/diluted in PBS to their final concentrations.

**Animal models**

Sprague Dawley rats were used for functional corneal healing experiments and wound-healing assessments. For mechanistic and molecular studies, genetically modified mouse lines were used, including B6.129P2(SJL)-Ntrk1tm1Ddg/J (TrkA^F592A^), Tg(KRT14-cre/ERT)20Efu/J (Krt-14-Cre), CBA;B6-Tg(Sox10-iCre/ERT2)388Wdr/J (Sox10-iCreERT2), B6;129S6-Gt(ROSA)26Sortm14(CAG-tdTomato)Hze/J (R26-LSL-tdTomato), and B6;129-Gt(ROSA)26Sortm1(DTA)Mrc/J (R26-LSL-DTA). The p75flox/flox mouse line (*63*) was generously shared by Brian Pierchala PhD (Indiana University School of Medicine). All transgenic lines were maintained on a C57BL/6J background.

Animals were housed in a temperature- and humidity-controlled environment under a 12-hour light/dark cycle with free access to food and water. All surgical procedures were performed under inhalational anesthesia using 2% isoflurane in oxygen in an aseptic manner. Meloxicam (4 mg/kg) was administered preoperatively and every 72 hours postoperatively for analgesia. All animal experiments were approved by the Indiana University Institutional Animal Care and Use Committee and were conducted in accordance with the ARVO Statement for the Use of Animals in Ophthalmic and Vision Research.

**Corneal denervation**

A stereotactic corneal denervation model was used to induce neurotrophic keratopathy (NK) in rodents, as previously described (*48, 92*)**.** Rats with an age of 250–300 g were used for functional studies, whereas mice with an age of 30–50 g were used for mechanistic experiments. Briefly, animals were anesthetized and positioned in a stereotactic frame. After a midline cranial incision was made and bregma identified, a small burr hole was created using stereotactic coordinates targeting the ophthalmomaxillary branch of the trigeminal nerve. Coordinates for rats were anterior-posterior (AP) 1.57 mm and mediolateral (ML) 1.87 mm, whereas coordinates for mice were AP 2.30 mm and ML 1.53 mm.

An insulated monopolar electrode (SonoPlex II Facet S; Pajunk GmbH, Geisingen, Germany) with the distal tip exposed (1 mm for rats and 0.5 mm for mice) was gently advanced through the burr hole until resistance from the skull base was encountered. Electrocautery ablation was then performed using an electrosurgical generator (Force FC-8C; Medtronic, Fridley, MN, USA) at 10 W for 60 seconds in rats and 6 W for 30 seconds in mice. Successful denervation was confirmed by loss of the blink reflex in response to cold saline stimulation of the affected eye. Following the procedure, the cranial incision was sutured and temporary tarsorrhaphy was performed, typically on the left eye.

**Corneal de-epithelialization**

Five days after corneal denervation, standardized corneal epithelial injury was performed under anesthesia The corneal epithelium was carefully removed using an Amoils rotating brush (Innovative Excimer Solutions, Toronto, ON, Canada). The rotating brush was lightly applied to the central cornea for approximately 5 seconds to create a uniform epithelial defect. During the procedure, cornea was gently stabilized using curved forceps fitted with silicone tubing to minimize unintended injury to the stroma and the limbal epithelium.

**Corneal healing assay**

Following corneal de-epithelialization in innervated or denervated eyes, with or without treatment, with NGF (50 µg/ml), αNGF (50 ng/ml), proNGF (50 µg/ml), αproNGF,(0.7 ug/ml), THX-B (5 µg/ul), MIM-D3 (25 mM), and LM11A-31 (25 µg/ul), topical treatments were administered under inhalational anesthesia for 10 min per treatment at approximately 10 µL per eye. In mouse model TrkA^F592A^ for selective TrkA inhibition, 1NMPP1 was administered through intraperitoneal injection (100 ul of 20 uM solution). The initial epithelial defect was visualized using fluorescein sodium staining (I-GLO Fluorescein Sodium ophthalmic strips; JorVet, Loveland, CO, USA). Briefly, approximately 10 µL of fluorescein sodium was applied topically to the corneal surface for 30 seconds, followed by gentle PBS washes to remove excess dye. Corneal epithelial healing was subsequently monitored daily after anesthetizing the animals and temporarily opening the tarsorrhaphy.

Fluorescence images were captured immediately after injury and every 24 hours thereafter for up to 96–120 hours using a Nikon D5100 digital camera (Nikon, Tokyo, Japan). Residual fluorescein-stained areas were used to assess wound closure and epithelial healing over time. Following imaging, the denervated eye was carefully reclosed with tarsorrhaphy until the next evaluation time point.

**AI-based corneal opacity grading**

To quantitatively assess the level of rodent corneal opacification following exposure to experimental conditions, a convolutional neural network (CNN) was developed using EfficientNet-B2, a deep learning architecture pretrained on the ImageNet dataset and fine-tuned for this task. The model used eye photographs as input and generated an opacity grade from 0 to 4, where 0 indicated a clear cornea and 4 indicated dense opacity.

The training dataset was generated from a blinded survey of laboratory members who independently graded randomly presented eye images for corneal opacity. The dataset included 55 images and 550 individual opacity scores. Each reviewer’s score was treated as an independent training example rather than averaging scores across reviewers, allowing the model to capture natural variability in human grading.

Model performance was evaluated using 5-fold cross-validation. Agreement between predicted and human-assigned grades was assessed using quadratic weighted kappa, an ordinal grading metric that accounts for the magnitude of disagreement between scores. The model achieved a quadratic weighted kappa of 0.851, indicating strong agreement with human graders.

**Schwann cell visualization and ablation**

To induce Cre-mediated recombination in corneal Schwann cells, tamoxifen-containing ophthalmic ointment was prepared by heating Systane eye ointment (Alcon, Geneva, Switzerland) to 60°C until liquefied, followed by addition of tamoxifen powder (T5648; Sigma-Aldrich, St. Louis, MO, USA) at a concentration of 25 mg/ml. The mixture was maintained at 60°C in the incubator (SciGene, California, USA) set to 5-10 rpm rotation for 2-3 hours until the tamoxifen was completely dissolved. Prepared ointment was protected from light, stored at 4°C, and used within 3 days of preparation.

For Cre activation, tamoxifen-containing ointment was applied topically to the corneal surface under isoflurane anesthesia for approximately 10 minutes daily for 2 consecutive days. In vivo imaging of tdTomato-positive Schwann cells within the cornea was then performed using a ZEISS Axio Zoom.V16 microscope (ZEISS Group, Oberkochen, Germany) while animals were maintained under 2% isoflurane anesthesia in a stereotactic frame.

**LESC Culture**

***Feeder cells***

NIH-3T3 fibroblasts were purchased from the American Type Culture Collection (ATCC; Manassas, VA, USA) and maintained in Dulbecco’s Modified Eagle Medium (DMEM) supplemented with 10% fetal bovine serum (FBS) and 1% HyClone Antibiotic Antimycotic. At approximately 70% confluence, feeder cells were treated with 4 µg/mL mitomycin C (Sigma-Aldrich, St. Louis, MO, USA) for 2 h at 37°C in a humidified incubator with 5% CO₂ to induce growth arrest. Cells were then washed, trypsinized, and seeded at a density of 4.5 × 10⁴ cells/cm².

***Limbal epithelial cell culture***

Primary human LESCs were isolated from cadaveric donor eyes of four different subjects obtained from VisionFirst Eye Bank (Indiana, USA). Donor eyes were used within a maximum of 5 days post-mortem. LESCs isolation was performed as previously described (*3*). Briefly, eyes stored in Optisol medium were transferred to Petri dishes containing 2.4 U/mL Dispase II (Gibco, Life Technologies, USA) and incubated at 4°C for 12 h. The epithelial sheet was then gently separated and incubated in 0.25% trypsin–0.02% EDTA for 30 min at room temperature to obtain a single-cell suspension.

Isolated cells were cultured in co-culture with mitomycin C–treated, growth-arrested NIH-3T3 feeder cells using Green’s medium consisting of 60% DMEM, 30% DMEM/F12, and 10% fetal calf serum (FCII), supplemented with 1 mM L-glutamine, 1 mM sodium pyruvate, 0.2 mM adenine, 5 µg/mL insulin, 0.5 µg/mL hydrocortisone, 10 nM cholera toxin, and 10 ng/mL epidermal growth factor (EGF). Cells were passaged at approximately 80% confluence using TrypLE Express (Thermo Fisher Scientific) and used for experiments up to passage 3 at 70–80% confluence.

***Colony-forming efficiency assay***

For colony formation, mitomycin C–treated NIH-3T3 feeder cells were seeded at a density of 2.6 × 10⁴ cells/cm². The following day, LESCs were plated onto the feeder layer at a density of 100 cells/cm² and maintained in green media for 7–14 days, with medium changes every 48 h. Where indicated, NGF (50 ng/ml), αNGF (50 ng/ml), proNGF (50 ng/ml), THX-B (5ng/ml), and MIM-D3 (50nM) were added and maintained throughout the duration of the experiment. Upon the appearance of well-defined colonies, cultures were fixed with 10% formalin and stained with rhodamine B (ThermoFisher Scientific, Waltham, MA). Colony images were acquired using a surgical light microscope (Leica, Wetzlar, Germany). Quantification was performed using ImageJ software, and clonogenic capacity was expressed as colony-forming efficiency (CFE), determined by the ratio of the mean intensity relative to the number of LECs initially seeded.

**Immunofluorescence staining for Biomarkers**

To confirm LESCs' identity and receptor expression, cells were immunostained for ΔNp63 and keratin 15 (K15) and co-stained for p75NTR and TrkA. To maintain monolayer cultures without feeder layers, cells were cultured in serum-free complete corneal epithelial cell basal medium (ATCC, Manassas, VA, USA) at 37°C in a humidified incubator with 5% CO₂. This medium contains essential and non-essential amino acids, vitamins, organic compounds, trace minerals, and inorganic salts and was supplemented with a corneal epithelial cell–specific growth kit (ATCC), including apo-transferrin, epinephrine, extract P, and hydrocortisone hemisuccinate. This complete media system supports corneal epithelial cell growth without the use of feeder layers, extracellular matrix proteins, or additional substrates. When adherent cells reached approximately 90% confluence, they were fixed with 4% formalin for 30 min at room temperature, PBS, and permeabilized with 0.05% Triton X-100 for 30 min. Following PBS washes, cells were blocked with 5% normal donkey serum (Jackson ImmunoResearch, West Grove, PA, USA) for 1 h at room temperature. Cells were incubated overnight at 4°C with the following primary antibodies: TP63 (1:100; NB110-59918, Novus Biologicals), Krt15 (1:100; Santa Cruz Biotechnology), p75NTR (1:100; Cell Signaling Technology), and TrkA (1:100; sc-80398, Santa Cruz Biotechnology). After washing with PBS, cells were incubated for 1 hour at room temperature with species-appropriate Alexa Fluor–conjugated secondary antibodies (Alexa Fluor 488, Cy5, or Cy3; 1:100; Jackson ImmunoResearch). Nuclei were counterstained with DAPI (1:2000; Thermo Fisher Scientific), and coverslips were mounted using Fluoromount-G mounting medium (Thermo Fisher Scientific). Images were acquired using a Zeiss Axio Imager microscope equipped with an ApoTome 3 system (Carl Zeiss, Germany).

**Whole-mount cornea staining**

Mouse or rat corneas were harvested, rinsed in PBS, and immediately fixed for 1 hour at 4°C in Zamboni’s fixative (4% paraformaldehyde and 15% saturated picric acid). Tissues were subsequently washed in PBS and permeabilized with ice-cold methanol at −20°C for 30 min. After additional PBS washes, samples were blocked with 5% NDS in PBS containing 0.1% Tween-20 and incubated overnight at 4°C with the following primary antibodies: βIII-tubulin (TUBB3; 1:100; #801201, BioLegend), TP63 (1:100; NB110-59918, Novus Biologicals), keratin 15 (Krt15; 1:100; Santa Cruz Biotechnology), p75^NTR^ neurotrophin receptor (p75^NTR^; 1:100; Cell Signaling Technology), and TrkA (1:100; sc-80398, Santa Cruz Biotechnology). Following PBS washes, corneas were incubated for 1 h at room temperature with species-appropriate Alexa Fluor–conjugated secondary antibodies (Alexa Fluor 488, Cy3, or Cy5; 1:100; Jackson ImmunoResearch). Nuclei were counterstained with DAPI (1:2000; Thermo Fisher Scientific), and tissues were mounted using Fluoromount-G mounting medium (Thermo Fisher Scientific). Images were acquired using a Zeiss Axio Imager microscope equipped with an ApoTome 3 system (Carl Zeiss, Germany). For epithelial nerve fiber quantification, extended depth-of-focus images were analyzed using the NeuronJ plugin in ImageJ software, and nerve fiber density was calculated as total nerve length per unit area.

**RNA isolation and Quantitative Real-time PCR**

For rat and mouse experiments, corneas were collected 5 days after corneal denervation and following three days of twice-daily pharmacological treatment. For cell culture experiments, treatments were added to the culture medium at the time of cell seeding, and cells were harvested on day 3. All treatment concentrations were used as described above appropriately. Enucleated corneas and cultured cells were immediately snap-frozen and stored until further processing. Total RNA was isolated from mouse corneas or cells using the miRNeasy Mini Kit (Qiagen, Düsseldorf, Germany) according to the manufacturer’s instructions. RNA concentration and purity were assessed using Nanodrop. Complementary DNA (cDNA) was synthesized from total RNA using the Verso cDNA Synthesis Kit (Thermo Fisher Scientific, Waltham, MA, USA) following the manufacturer’s protocol. Quantitative real-time PCR (RT-qPCR) was performed using SYBR Green chemistry on a QuantStudio 3 Real-Time PCR System (Applied Biosystems, Alameda, CA, USA). Amplification was carried out using standard cycling conditions as recommended by the manufacturer. Relative gene expression levels were calculated using the ΔΔCt method and normalized to the GAPDH housekeeping gene. Primer sequences used for amplification are listed in Table below.

| Gene | Forward | Reverse |
| --- | --- | --- |
| ABCG2 | CAGGGTCATTCAAGAGTTAGGTC | AGAACAAGATGGAAGGATCAGTG |
| KRT15 | TGCGACTACAGCCAATACTTC | GGCCAGCTCATTCTCATACTTG |
| P63 | TTCGGACAGTACAAAGAACGG | GCATTTCATAAGTCTCACGGC |
| KRT3 | GAGCGGGAACAGATCAAGAC | TGCCTGAGATGGAACTTGTG |
| KRT12 | CCAAATCACAAGCACAGTCAAC | CTCACCATTCACCATCTCCTG |
| GPHA2 | TCCCTTCTCGGTACTCTGTG | ACTGCAGCTGTACTTTGACC |
| KI67 | AAAAGAATTGAACCTGCGGAAG | AGTCTTATTTTGGCGTCTGGAG |
| GAPDH | ACATCGCTCAGACACCATG | TGTAGTTGAGGTCAATGAAGGG |
| Abcg2 | ACGACTGGTTTGGACTCAAG | AGTAAGGTGAGGCTGTCAAAC |
| Krt15 | GAGATGAGGGAGCAGTATGAAG | CTGTGATCTCCGTCTTGCTG |
| P63 | TGCCATGCCTGTCTACAAG | GCTGTTCCCTTCTACTCGAATC |
| Krt12 | ATCCCTTTCTGTCTCCAAACC | ACTTCGCCATTCACTATCTCC |
| Gpha2 | CACAACATCACCTCTTCCTCC | ACGGCACATATCACACTGG |
| Ki67 | TGCCCGACCCTACAAAATG | GAGCCTGTATCACTCATCTGC |
| Gapdh | CTTTGTCAAGCTCATTTCCTGG | TCTTGCTCAGTGTCCTTGC |

**Western blot analysis**

For corneal denervation experiments, rodent corneas were collected 5 days after denervation. For pharmacological treatment experiments, corneas were collected following 3 days of twice-daily treatment. For de-epithelialization studies, corneas were harvested at 5, 15, and 60 minutes after the procedure. All treatments were the same concentrations used in the corneal epithelial healing assays, with MIM-D3 administered at 25 μM. Isolated mouse and rat corneas were immediately snap-frozen, and the epithelial layer was gently scraped to collect epithelial cells in ice-cold radioimmunoprecipitation assay (RIPA) buffer containing 25mM Tris-HCl (pH 7.6), 150 mM NaCl, 1% NP-40, 1% sodium deoxycholate, 0.1% SDS, and 5mM EDTA, supplemented freshly with protease and phosphatase inhibitor cocktails (Roche, Basel, Switzerland) before use. Lysates were then clarified by centrifugation at 12,000 × g for 15 min at 4°C, and protein concentrations were determined using the Pierce Bradford Plus Protein Assay Kit (A55866; Thermo Fisher Scientific) according to the manufacturer’s instructions. Equal amounts of protein (40 µg per lane) were resolved on pre-cast SDS–polyacrylamide gels (Bio-Rad) and transferred to polyvinylidene difluoride (PVDF) membranes (Bio-Rad). Membranes were blocked with 2% bovine serum albumin (BSA; Thermo Fisher Scientific) in Tris-buffered saline (TBS) for 1 h at room temperature and incubated overnight at 4°C with the following primary antibodies: phospho-TrkA (Y490; 1:1000; Cell Signaling Technology, #9141), total TrkA (1:1000; Invitrogen, PA5-109216), p75NTR (1:1000; EMD Millipore, #07-476), phospho-JNK (Thr183/Tyr185; 1:1000; Cell Signaling Technology, #9251), total JNK (1:1000; Cell Signaling Technology, #9252), phospho-AKT (Thr308; 1:1000; Cell Signaling Technology, #9275), total AKT (1:1000; Cell Signaling Technology, #9272), and total ERK1/2 (1:1000; Cell Signaling Technology, #9102). Following washing with Tris-buffered saline containing 0.05% Tween-20 (TBST), membranes were incubated with horseradish peroxidase–conjugated goat anti-rabbit IgG secondary antibody (1:5000; Cell signaling technology) for 1 hour at room temperature. Protein bands were visualized using enhanced chemiluminescence (ECL) reagents and detected using a digital imaging system. Band intensities were quantified using ImageJ software, and protein activation was calculated as the ratio of phosphorylated protein to total protein.

**Single-cell RNA sequencing**

Previously published single-cell RNA sequencing data from innervated and denervated rat limbal tissues (*12*) were reanalyzed using the Seurat R package. Following standard quality control to remove cells with low gene counts or high mitochondrial expression, the datasets were normalized, scaled, and integrated using an anchor-based workflow to correct for batch effects. Dimensionality reduction was performed via Principal Component Analysis (PCA), followed by SNN graph-based clustering (resolution 0.5) and UMAP visualization, with cell clusters annotated based on established marker genes.

**SUPPLEMENTARY FIGURES**

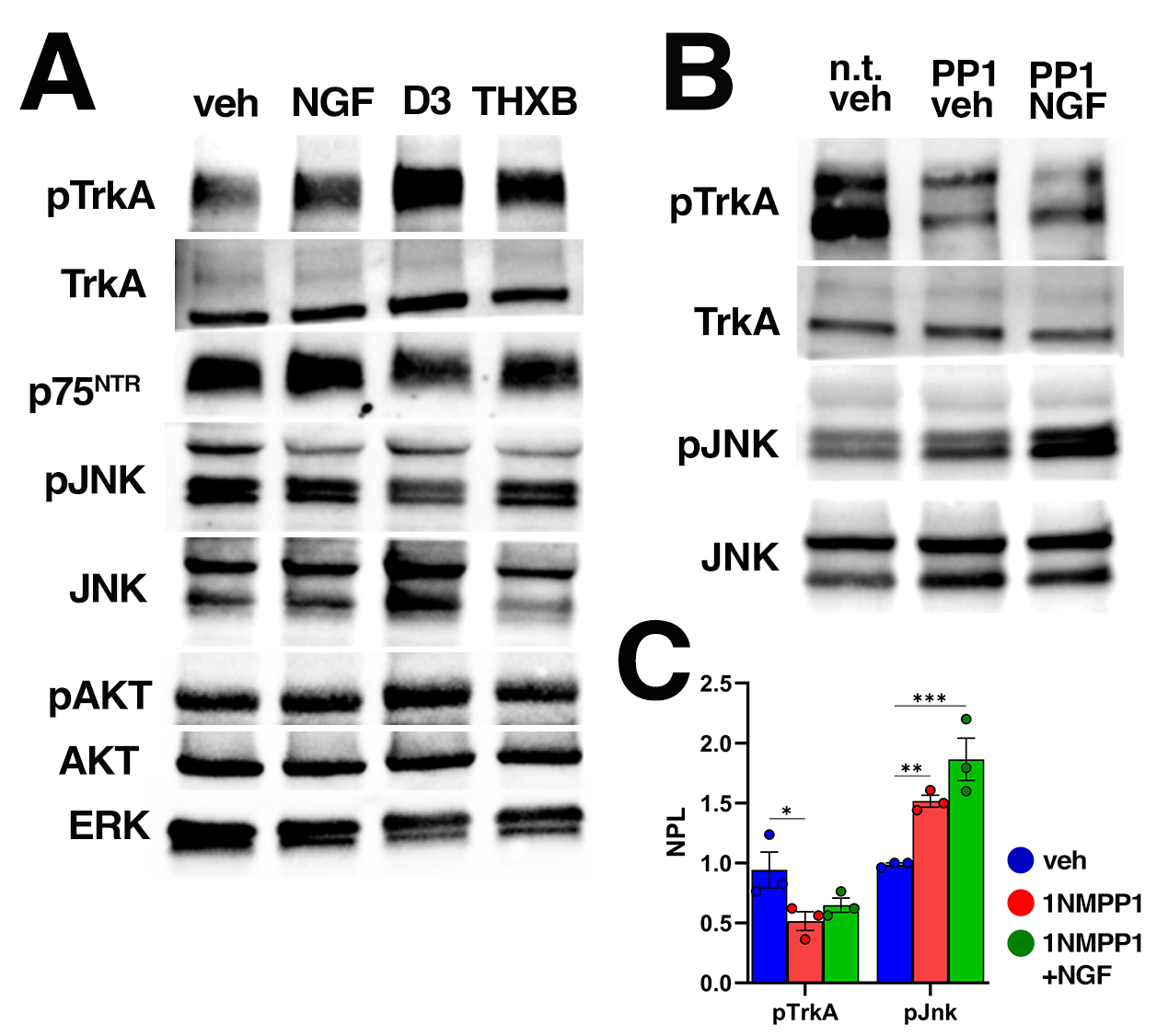

**Figure S1.** (**A**), (**B**) Western blot analysis of corneal epithelial lysates, demonstrating the expression/phosphorylation of the indicated proteins in healthy rat in (A) or TrkA^F592A^ mouse in (B) corneas exposed to every of the indicated topical compounds. In (A) – three consecutive days of two daily topical administrations. In (B) – three daily intraperitoneal injections with 1NMPP1 (PP1) or PBS (non-treated “n.t”) along with two daily topical rhNGF or vehicle (“veh”) alone. (**C**) Quantitative representation of normalized protein level (“NPL”) in (B). *n* = 3, ANOVA.

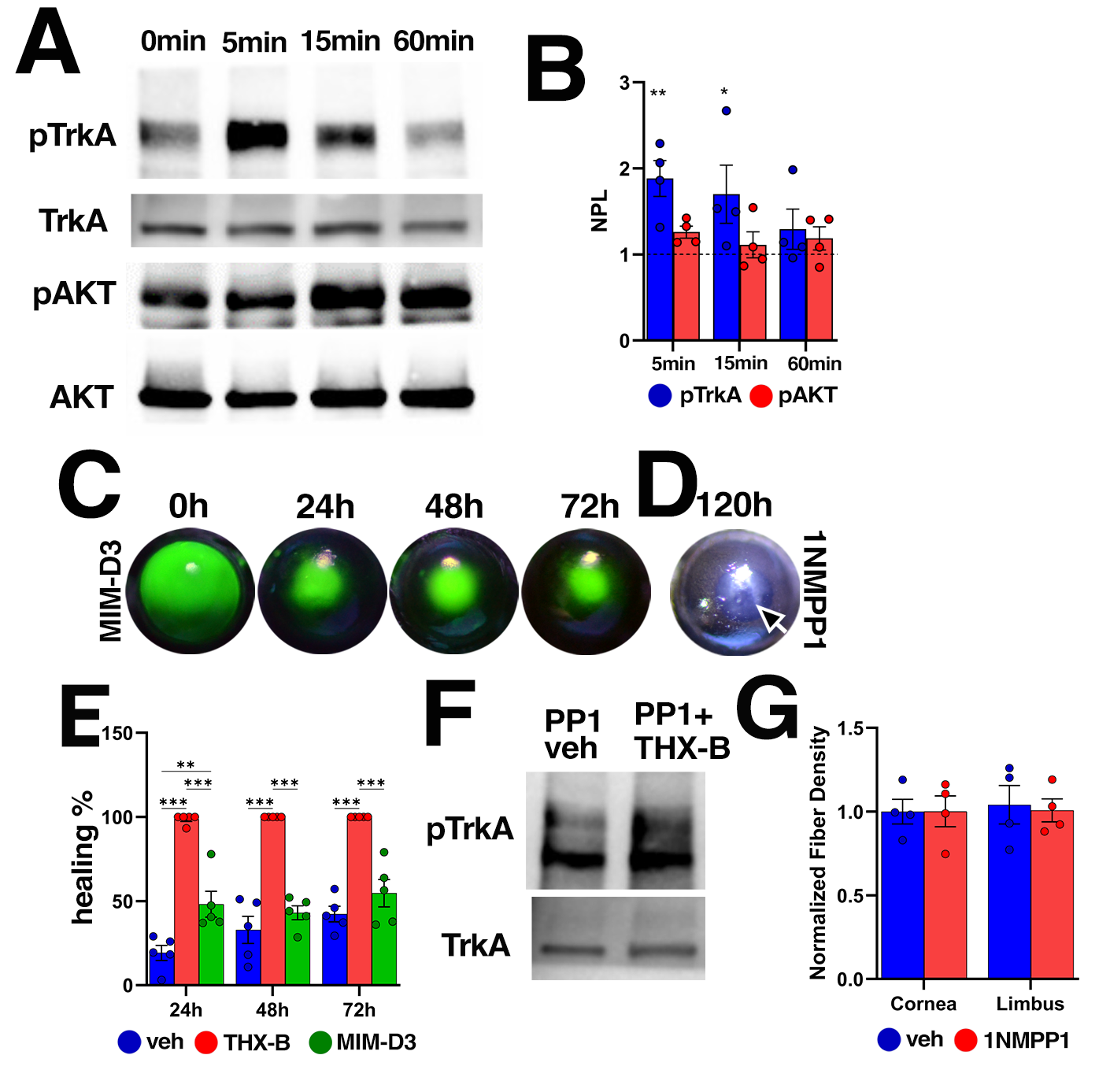

**Figure S2.** (**A**) Western blot analysis of corneal epithelial lysates, demonstrating the expression/phosphorylation of TrkA and AKT at 5, 15 and 60 minutes post de-epithelialized corneas. (**B**) Quantitative representation of normalized protein level (“NPL”) in (A). The punctate line at “1” represents base line proteins phosphorylation of non-wounded cornea. n= 4, ANOVA. (**C**) Representative live photos of experimentally de-epithelialized 1NMPP1 pre-treated TrkA^F592A^ and topical tavilermide (MIM-D3) treated mouse corneas as in Figure 3C,D, demonstrating the course of the epithelial healing as per the fluorescein staining. (**D**) Representative live bright field images 120 hours post de-epithelialization, as per (C). (**E**) Quantitative representation of the course of the wound healing in presence of topical MIM-D3, as in (C), or vehicle treated alone or THX-B (Figure 3K) of TrkA-inhibited de-epithelialized corneas. *n* ≥ 5, ANOVA. (**F**) Representative western blot analysis of corneal epithelial lysates from TrkA^F592A^ mice receiving three consecutive daily intraperitoneal injections of 1NMPP1 and two daily topical administrations of vehicle alone (“veh”) or THX-B, demonstrating TrkA phosphorylation status under each condition. (**G**) Quantitative representation of the immunostaining results in Figure 3I, demonstrating comparable limbal and corneal innervation in untreated or 1NMPP1 pre-treated TrkA^F592A^ mice corneas. *n* = 4, ANOVA.

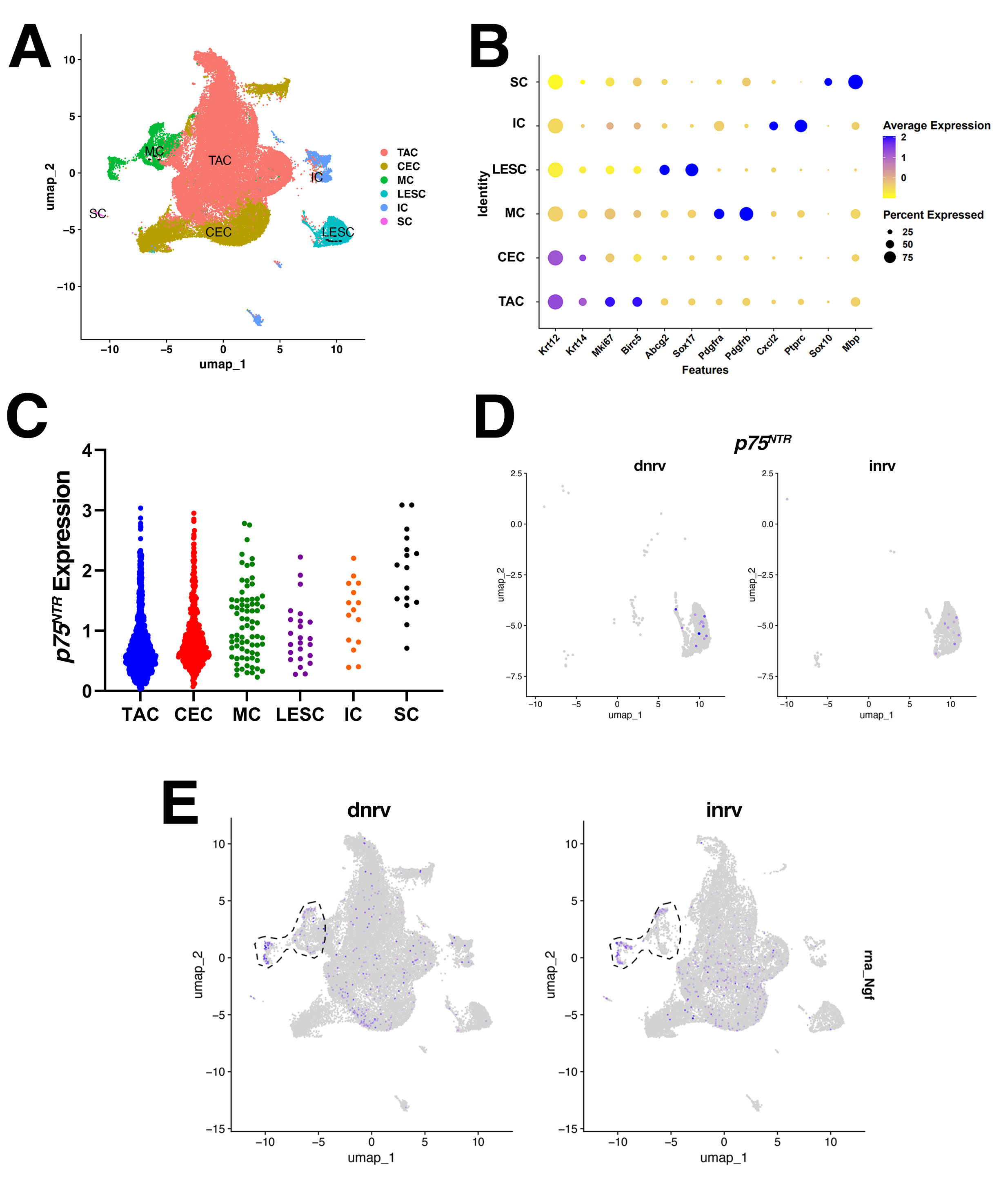

**Figure S3.** Analysis of the published scRNA-seq data of the limbal area of healthy, denervated or centrally de-epithelialized rat corneas (*12*). (**A**) UMAP the indicated corneal epithelial cell clusters. TAC – transient amplifying cells, CEC – corneal epithelial cells, MC – mesenchymal cells, LESC – limbal epithelial stem cells, IC – immune cells, SC – Schwann cells. (**B**) Dot plot representation of the indicated genes scRNA-seq expression distribution across the indicated cell clusters. (**C**) Violin plot showing relative corneal epithelial expression of the *p75^NTR^* (*ngfr*) mRNA by the indicated cell clusters. (**D**) Qualitative comparative representation of *ngfr* mRNA expression by the LESC cluster in innervated (inrv) and denervated (dnrv) cornea. (**E**) Qualitative comparative representation of *ngf* mRNA expression by the mesenchymal cell cluster in innervated (inrv) and denervated (dnrv) cornea.

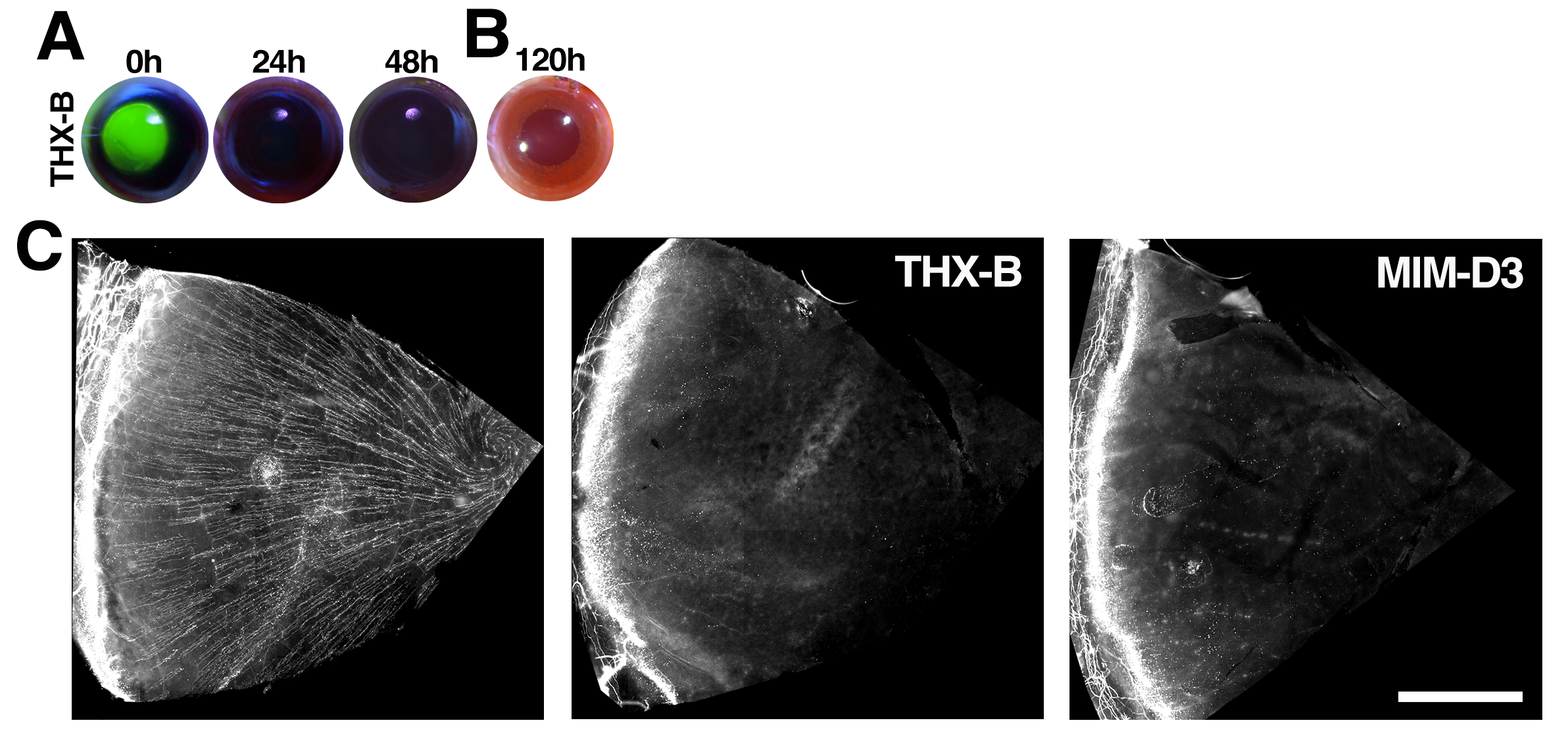

**Figure S4.** (**A**) Representative live photographs of fluorescein-stained rat normally innervated de-epithelialized corneas, demonstrating the course of corneal epithelial healing in response to topical treatment with THX-B. **0h** indicates corneal immediately after de-epithelialization. (**B**) Representative live bright field images of the corneas as per (A) 120 hours after de-epithelialization. (**C**) Representative immunofluorescent 15μm extended-depth focus images of whole mount βIII-tubulin-labeled normally innervated (left image) or denervated, topically treated as in with THX-B or tavilermide (MIM-D3) (as per Figure 5A) rat quoter corneas, demonstrate complete ablation of the corneal nerves in the denervated corneas. Scale bar – 1mm.

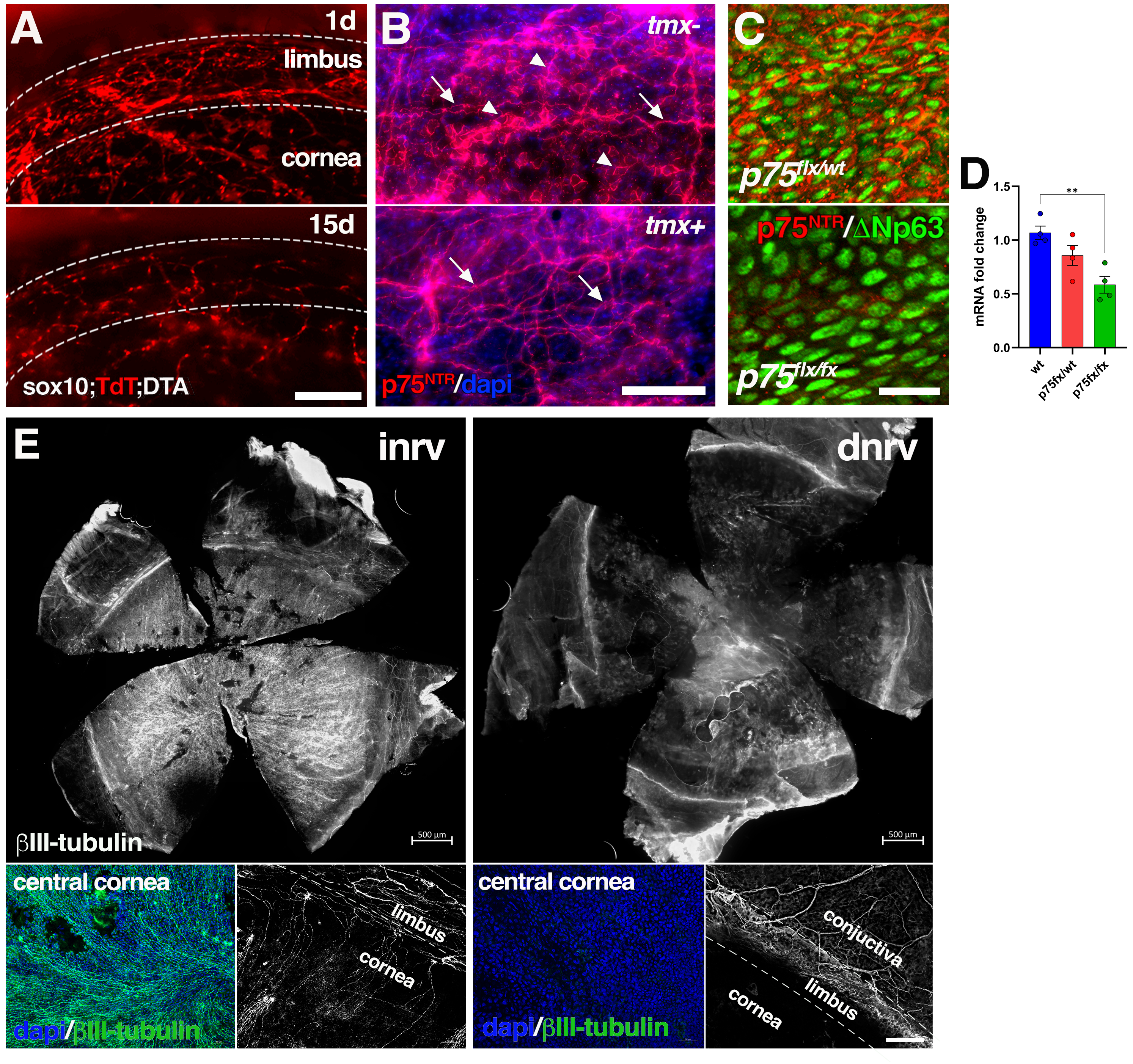

**Figure S5.** (**A**) Representative live image of Sox10-TdT;DTA mouse cornea at the day one (upper image) and the day 15 (lower image) since the first, out of the two daily, topical tamoxifen administration shows nearly complete ablation of the TdT-positive Schwann cells (red) in the limbal area of the fifteen days post-Cre-activated cornea. Scale bars - 250µm. (**B**) Immunofluorescent images of the limbal area of homozygous *K14-p75^fx/fx^* demonstrate presence of both p75^NTR^-positive (red) nerves (arrows) and sub-epithelial cell membrane (arrowheads) in tamoxifen-non-treated cornea (upper image), but only presence of p75^NTR^-positive nerves but absence of p75^NTR^-positive cells in tamoxifen-treated cornea (lower image). Scale bar – 100μm. (**C**) Immunofluorescent images of the limbal area of homozygous *K14-p75^fx/fx^* (lower image) and heterozygous *K14-p75^fx/wt^* (upper image) tamoxifen treated mouse corneas demonstrate presence of p75^NTR^-positive (red; upper) and p75^NTR^-negative (lower), and nuclear ΔNp63-positive (green) cells. Scale bar - 25μm. (**D**) RT-qPCR analysis of p75^NTR^ mRNA in the corneal epithelium of *wt, K14-p75^NTR fx/wt^* or *K14-p75^NTR fx/fx^* tamoxifen treated mice, demonstrates gradual reduction of total p75^NTR^ transcription following induced heterozygous or homozygous LESC/TAC-specific gene deletion. The corneas were harvested after two constitutive daily topical tamoxifen administration (*12*). n= 4, ANOVA. (**E**) Representative immunofluorescent images of whole mount βIII-tubulin-labeled normally innervated (left panel) or denervated (right panels) of p75^NTR^ null mouse cornea, harvested five days after denervation. The images at lower panels of the central cornea (merged with dapi, left) or βIII-tubulin alone (right) are representative magnified areas of the central and the limbal cornea, respectively, as at the upper panels, demonstrate complete absence of axons in the denervated cornea. Scale bars: upper panel – 500 μm; lower panel – 100 µm.

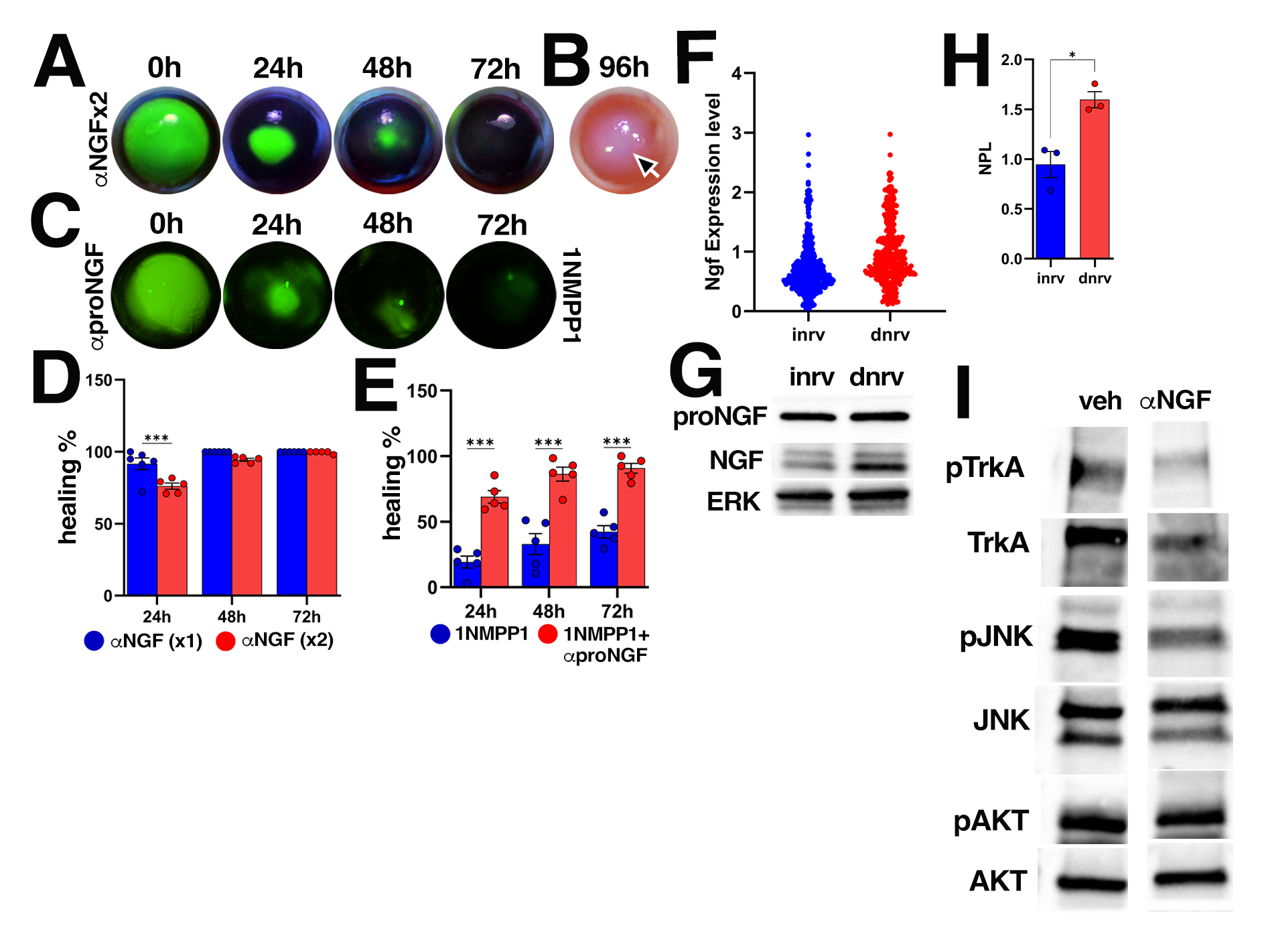
**Figure S6.** (**A**), (**C**) Representative live photographs of fluorescein-stained rat in (A) or 1NMPP1 pre-treated (as in Figure 3C) TrkA^F592A^ mouse in (C), normally innervated de-epithelialized corneas, demonstrating the course of epithelial healing in response to topical two daily of αNGF (50 ng/ml) in (A), or one daily of αproNGF (0.7 μg/ml) in (C) treatment, started at the day of de-epithelialization. (**B**) Representative live bright field image of the corneas as per (A). (**D**), (**E**) Quantitative representation of the results in (A), (C), respectively, in comparison to one daily αNGF (as in Figure 6C,E) in (**D)** or 1NMPP1 pre-treated and topical vehicle alone treated (as in Figure 3C,D,K) corneas. (**F**) Violin plot showing relative total corneal epithelial expression of the *NGF* mRNA in innervated (“inrv”) or denervated (“dnrv”) corneas. (**G**) Western blot analysis of rat corneal epithelial lysates, comparing proNGF and NGF protein expression between innervated and denervated corneas. (**H**) Quantitative representation of NGF normalized protein level (NPL) as per (G). (**I**) Representative western blot analysis of epithelial lysates derived from untreated or topical αNGF (50 ng/ml; two daily doses for three constitutive days) treated rat corneas demonstrating comparable AKT but reduced JNK phosphorylation in response to αNGF treatment.
